# Navigational Frames of Reference as Critical Regulators of Hippocampal Interneuron Coding Properties

**DOI:** 10.1101/2025.02.07.637101

**Authors:** Jean-Bastien Bott, Lorene Penazzi, Salodin Al-Achkar, Minyoung Cho, Bruno Rivard, Etienne Gautier Lafreniere, Sylvain Williams

## Abstract

Efficient spatial navigation relies on the hippocampus integrating local (proximal) and global (distal) cues, collectively called frames of reference, to guide behavior and support memory. Although these cues control the anchoring of principal cell fields, how these frames tune interneuron functions remains unknown. Traditionally, interneurons such as O-LM and VIP cells have been viewed primarily as speed encoders, although some also encode spatial information or respond to discrete stimuli. Using calcium imaging in freely behaving mice performing a new spatial learning task that differentiates between reference frames, we demonstrate that O-LM cells displayed a striking bimodal activity pattern, altering both their speed and spatial encoding properties. In contrast, VIP interneurons were largely unaffected by changes in the frame of reference, instead correlating with familiarization. Notably, linear decoding using speed scores revealed that only O-LM interneurons provide an accurate readout of the dominant reference frame, enabling prediction of the animal’s navigation strategy. These findings highlight that hippocampal interneurons can flexibly adapt their functions depending on cognitive factors such as the reference frames used to guide behavior.

## Main

Spatial navigation is a remarkable process that relies on the integration of local and distal cues, allowing organisms to orient and move effectively in their environment. To navigate successfully, individuals must continuously update their spatial position and heading based on local features such as environmental geometry and distal landmarks that often require active visual sampling^1^. These local and distal cues serve as reference frames that guide the activity of hippocampal principal neurons, especially place cells, which fire at specific locations and encode spatial information^2–6^. Changes in these environmental cues can markedly alter both the activity of principal neurons and the structure of spatial coding^7,8^. Initial studies underscored the predominance of distal cues in shaping place fields^2,9–14^. Later findings revealed that local environmental features also influence place cell firing, establishing spatial representations tied to the specific arrangement and properties of an experimental setup (e.g., the shape and walls of a maze^15–20^). Consequently, variations in these reference frames profoundly affect principal cell activity and the nature of spatial coding^7^. Interneurons are central to hippocampal network function by regulating network excitability, controlling spike timing, synchronizing neuronal ensembles^21,22^, and encoding running speed to align place cell activity with locomotor velocity^23^. Despite advances in understanding how principal neurons utilize different frames of reference, how interneurons integrate spatial reference frames is unknown.

Understanding how interneurons shape hippocampal function requires examining their unique roles in information encoding and how their activity adapts to cognitive demands and experimental conditions. In hippocampal circuits, interneurons firing correlate with diverse behavioral states, ranging from immobility to active locomotion and increasing evidence highlights their critical role in mediating speed encoding in the hippocampus^24,25^. Speed information is believed to dynamically update the hippocampal map as an animal move, ensuring precise adjustments in spatial representation in response to changes in velocity^26–30^. Until now, it was widely presumed that these signals offer a stable, rate-coded velocity input that remains consistent across environments and over time^31,32^, enabling interneurons to integrate locomotor and spatial information into a coherent navigational framework^25,32–35^. Among these interneurons, CA1 Oriens-Alveus Lacunosum Moleculare (O-LM) cells, a somatostatin-positive interneuron (SOM) subtype, are well-positioned to modulate spatial memory, sensory integration, and synaptic plasticity through dendritic modulation of CA3 and entorhinal inputs^36–39^. Several studies indicate that O-LM cell firing correlates positively with running speed^2,35,40–43^. However, one report^44^ found a negative correlation, implying that O-LM speed encoding could be more complex than previously recognized and may involve yet-to-be-identified mechanisms. Vasoactive intestinal peptide (VIP) interneurons are thought to finely regulate O-LM cells to adjust hippocampal network gain and plasticity, ultimately influencing place cell activity and spatial coding^45^. While VIP neurons tend to be more active during novel experiences and SOM cells in familiar contexts^35,43^, whether these patterns reflect distinct navigational strategies used in goal-directed behavior remains an open question. Given that O-LM interneuron recruitment can be conditional and task-dependent^38^, we hypothesize that O-LM cells exhibit a flexible, context-sensitive pattern of activity determined by strategies related to the frame of reference.

Dissecting how local and distal frames of reference influence spatial navigation is challenging, especially in complex mazes where these frames often overlap^7,46–51^. In simpler paradigms, however, the ability to consistently identify a starting location has been proposed as a critical factor for the selection of a distal frame of reference^52–55^. Although this stems from simple mazes, the same principle may apply to more complex, multi-route tasks such as the Star maze^56^. To explore this possibility, we manipulated the ability to localize the starting point in the Star maze to bias navigation toward either a local or a distal frame of reference.

We used miniscope calcium imaging in freely moving mice to investigate how O-LM and VIP interneurons respond under these conditions. In both the Star maze and simpler tasks, O-LM speed coding, spatial tuning, and event-specific activation were strongly reference-frame dependent, whereas VIP cells generally maintained a stable, familiarization-associated activity across reference frames. These findings highlight how interneurons can flexibly adapt their function to the cognitive demands and behavioral strategies, challenging the traditional view that interneurons’ functions are stable across environments.

## Results

### Effects of the start point identification on navigation strategy and memory performances

To determine how the dominant frame of reference (local *versus* distal) during navigation impacts the activity of O-LM and VIP interneurons in a spatial memory task, we developed a new version of the Star maze^56^. This paradigm provides a multi-route, multi-choice environment that requires flexible trajectories for optimal navigation toward a single target location. The Star maze device consisted of an inner pentagon track connected to five independent arms (see **Methods**, **Fig. 1a**). Mice were given 16 trials per day, over three consecutive days, to learn the location of a sucrose water reward in the target arm. In this version of the Star maze, mice started each trial from a new pseudo-randomly selected start arm, ensuring hippocampal-dependent spatial reference memory. Each trial was divided into 4 phases (**Fig. 1b**): (i) During the start arm (SA) location phase, mice were confined to a pseudo-randomly selected start arm, blocked by a door, for 10 seconds. During this phase, mice could localize the starting point of the upcoming trial. (ii) In the navigation phase, after opening the door, mice freely explored the maze until they found the target arm, which remained at the same spatial location across the three training days. As learning progressed, navigation improved from mostly indirect trajectories involving visits to non-target (errors) arms (see **Supplementary Fig. 1a-b**) to predominantly direct trajectories toward the target arm. (iii) During the reward consumption phase, upon reaching the target arm, mice consumed the sucrose water reward. (iv) Finally, after consuming the reward, mice had 10 seconds to observe and encode the distal environmental landmarks associated with the target location.

**Fig. 1:**
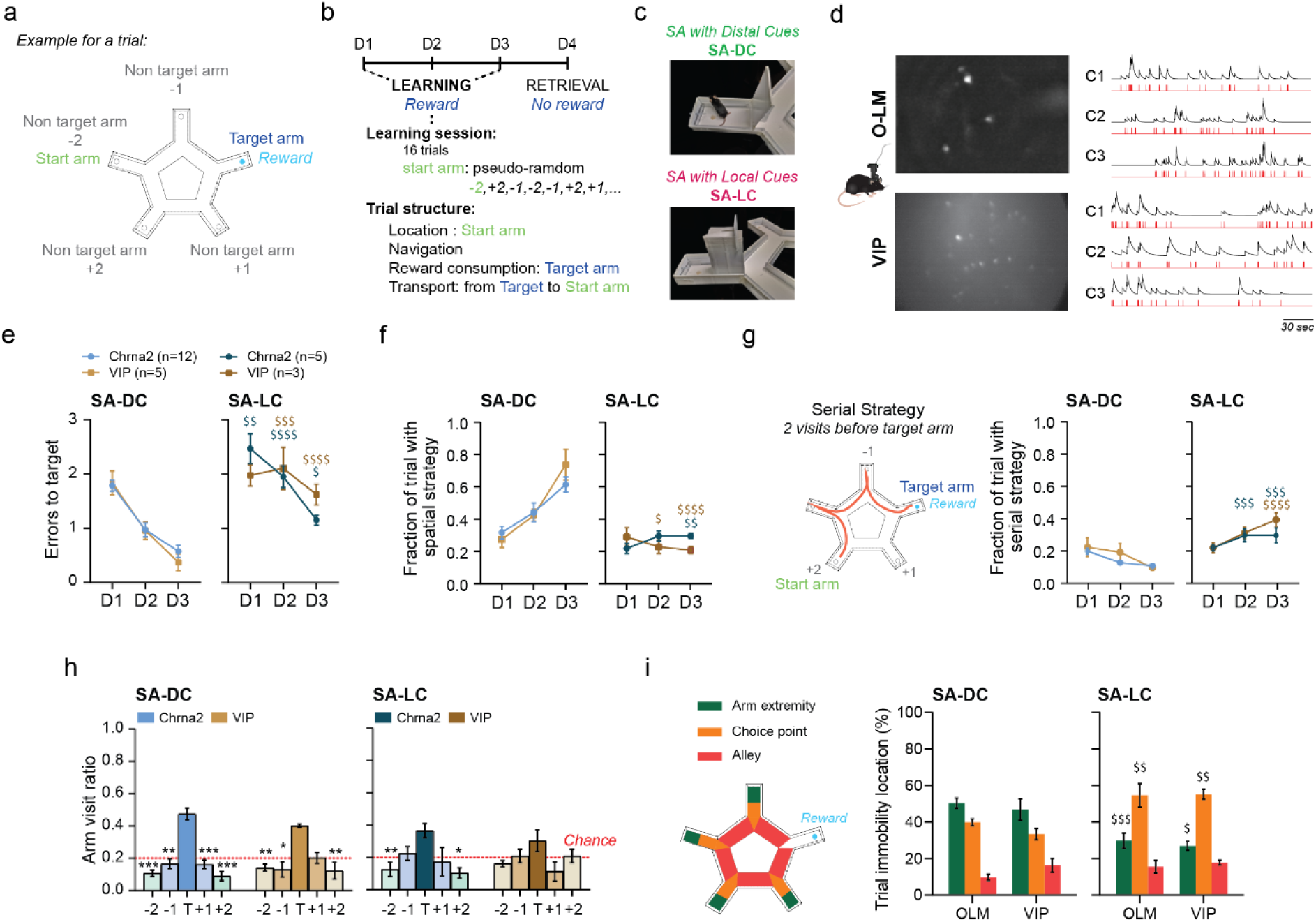
Distal visual cues accessibility during the start point location phase is crucial for optimal spatial reference memory. **a-b,** Schematic of the Star Maze apparatus. A water reward (light blue) is delivered at the TARGET arm (dark blue), which remains fixed over the three learning days. The four other arms are used as start arms (green), where mice are pseudo-randomly introduced and confined for 10 seconds (location period) before the navigation period begins with the opening of the door. Each learning day consists of 16 trials. **c,** Two conditions were used to manipulate visual access to distal landmarks during transportation and the location phase in the start-arm. In the SA-LC, animals are kept in a box during both transportation and the duration of the start arm location period. **d,** Calcium imaging was performed on freely moving mice using GCaMP6f. Left: DF/F maximum intensity projection of two representative fields of view illustrating the sparsity of O-LM interneurons compared to VIP interneurons in dorsal CA1. Scale bars, 50 µm. Right: Representative fluorescent signal (dark lines) and deconvoluted spike rates (red lines) from O-LM and VIP interneurons. **e-h,** Behavioral readouts showing the consequences of visual deprivation of distal cues during the location process: **e**, increased errors in the SA-LC condition for both genotypes (session * condition interaction F(2,60)=3.233, p=0.0464; genotype * condition * session interaction F(2,60)=2.590, p=0.0934; α=0.05). **f,** Preferential use of spatial strategies for both genotypes in the SA-DC condition (session * condition interaction F(2,15)=11.73, p=0.0009; genotype * condition * session interaction F(2,15)=2.307, p=0.1338; α=0.05). **g,** Left: example of a serial trajectory. Right: increased use of non-spatial strategies for both genotype in the SA-LC condition (session * condition interaction F(2,42)=12.85, p<0.0001; genotype * condition * session interaction F(2,42)=1.587, p=0.2166; α=0.05). **h,** A lower bias toward the target arm compared with non-target arms during memory retrieval in SA-LC condition (arm * condition interaction F(4,25)=4.194, p=0.0098; genotype * condition interaction F(1,25)=2.901e-011, p>0.9999; α=0.05). **i,** Left: regions used to classify immobility locations. Right: for both genotypes, mice from SA-DC condition were immobile preferentially at the arm extremities, while mice in the SA-LC condition immobility occurred preferentially at choice points (condition * location interaction: F(2,15)=26.22, p<0.0001; condition * location * genotype interaction: F(2,15)=0.5669, p=0.579, α=0.05). **Statistics: e-i,** 3-way ANOVA with LSD multiple comparisons post hoc test. Differences between conditions for the same genotype and measure $ p<0.05, $$ p<0.01, $$$ p<0.001 and $$$$ p<0.0001. **h,** Differ from non-target arms * p<0.05, ** p<0.01, *** p<0.001 and at **** p<0.0001. Data are shown as mean ± SEM.

Based on previous research emphasizing the importance of the starting point localization in navigation^52–55^, we hypothesized that in the Star maze accurate localization of the start arm is essential for establishing an optimized trajectory toward the target based on a distal frame of reference. Specifically, we predicted that preventing access to distal landmarks during the start-arm localization phase would disrupt spatial learning and memory performances. To test this, we designed two conditions. In the start arm with distal cues (SA-DC) condition, mice were transported to the start arm on a platform, allowing continuous visual access to distal landmarks throughout both the transportation and the subsequent waiting period (**Fig. 1c**). In contrast, in the start arm with only local cues (SA-LC) condition, mice were transported inside a closed and opaque box, remaining enclosed in this box during the waiting period, and therefore, could not view distal landmarks. (**Fig. 1c**). However, once the navigation phase began, both groups (SA-DC and SA-LC) had access to the exact same set of distal and local cues.

Analysis of the memory performances of mice implanted for further calcium imaging analysis (**Fig. 1d**) showed that, remarkedly, removing distal cue access only during the start arm location phase significantly impaired spatial learning. Under the SA-LC conditions, mice made more errors (i.e., visited non-target arms; **Fig. 1e**) and failed to improve their spatial trajectory efficiency, which remained at chance level. In contrast, mice in the SA-DC condition quickly adopted a spatial-based navigation strategy characterized by direct trajectories to the target, while in the SA-LC condition, mice persisted with non-spatial serial strategies (**Fig. 1f,g**). These results were similar in both Chrna2-cre and VIP-cre mice strains. Subsequent memory retrieval testing on Day 4 showed that mice from the SA-LC condition displayed poor target arm discrimination for both genotypes, suggesting weakened spatial memory (**Fig. 1h**). Even though mice from the SA-LC condition eventually had visual access to distal cues during the trial, they never achieved the same trajectory optimization as SA-DC controls. These finding highlights that even transient disruptions in access to distal landmarks at the beginning of the trajectory can have lasting effects on spatial navigation and memory.

Immobility periods during navigation may reflect cognitive processes such as planning, decision-making, and updating of spatial representations^57^. We hypothesized that the spatial distribution of these pauses would vary depending on whether navigation was guided by a distal or a local frame of reference. To explore this, we examined where immobility events occurred during the navigation phase in both Star maze paradigms. Under the SA-DC condition, as mice learned to navigate efficiently, immobility occurred predominantly at arm extremities (**Fig. 1i**), suggesting that these pauses, along with the start arm location phase, may facilitate distal landmark sampling and enhanced trajectory planning. In contrast, under the SA-LC condition, immobility events were concentrated at choice points reliant on local features such as walls and angles akin to a proximal serial strategy. These findings demonstrate that merely having distal cues available is not enough for efficient navigation; the animal must actively adopt a distal frame of reference to benefit from them. Indeed, when mice could not anchor their starting point under SA-LC conditions, they defaulted to a local frame of reference—even though the exact same distal cues were present as in SA-DC trials. This highlights that identifying the start location is crucial for engaging a distal frame of reference, ultimately enabling more effective allocentric navigation and stronger spatial memory. If hippocampal interneurons, such as O-LM and VIP cells, are sensitive to the chosen frame of reference, their activity patterns should reflect the animal’s navigational strategy and the accessibility of distal information.

### During the location phase, distal cues drive O-LM but not VIP activation

Building on our previous findings that the starting point location phase strongly dictates the selection of the reference frame and navigation strategy, we next examined how distal cue availability during this start-arm location phase affects O-LM and VIP interneuron activity. We selectively targeted O-LM interneurons using Chrna2-cre mice^36^ (see **Methods** and see **Supplementary Fig. 2c,d**). For miniscope calcium imaging, mice were implanted with a GRIN lens over the dorsal CA1 alveus surface to record activity from stratum oriens in Chrna2-cre mice and from stratum pyramidale in VIP-cre mice (**Fig. 1d**, and see **Supplementary Fig. 2**). We analyzed ΔF/F calcium activity aligned with the start arm’s beginning and ends, computing response reliability for each cell. In the SA-DC condition, where distal cues were accessible, most O-LM interneurons exhibited robust activation during the start arm, with the most pronounced activity occurring immediately upon the animal’s introduction to the start arm (**Fig. 2a**). In contrast, under the SA-LC condition, where distal cues were unavailable, O-LM interneuron activity was markedly reduced during the start arm phase. This striking difference indicates that O-LM cells depend highly on distal cues to activate during this critical location phase. VIP interneurons followed a different pattern: under SA-LC conditions, their initial activation on Day 1 was greater than in SA-DC, but this difference dissipated over the subsequent two days of training (**Fig. 2b**). This dynamic adjustment suggests that VIP interneurons may play a transient role in early learning, particularly when distal information is absent, but may be less critical for sustained spatial strategies. The SA-DC condition engaged a significantly larger proportion of O-LM cells than the SA-LC condition, reinforcing the notion that distal cues are essential for O-LM activation. In contrast, the proportion of VIP cells engaged during the start arm phase was only marginally influenced by distal cue availability (**Fig. 2c**). Interestingly, O-LM interneurons also displayed notable activity changes around the reward phase. In the SA-LC condition, O-LM cells exhibited a pronounced increase in activity approximately two seconds before reward consumption (**Fig. 2d-f**). This finding aligns with previous studies showing that somatostatin-positive (SOM) interneurons, likely including O-LM cells, are strongly modulated by reward signals^35,58^.

**Fig. 2:**
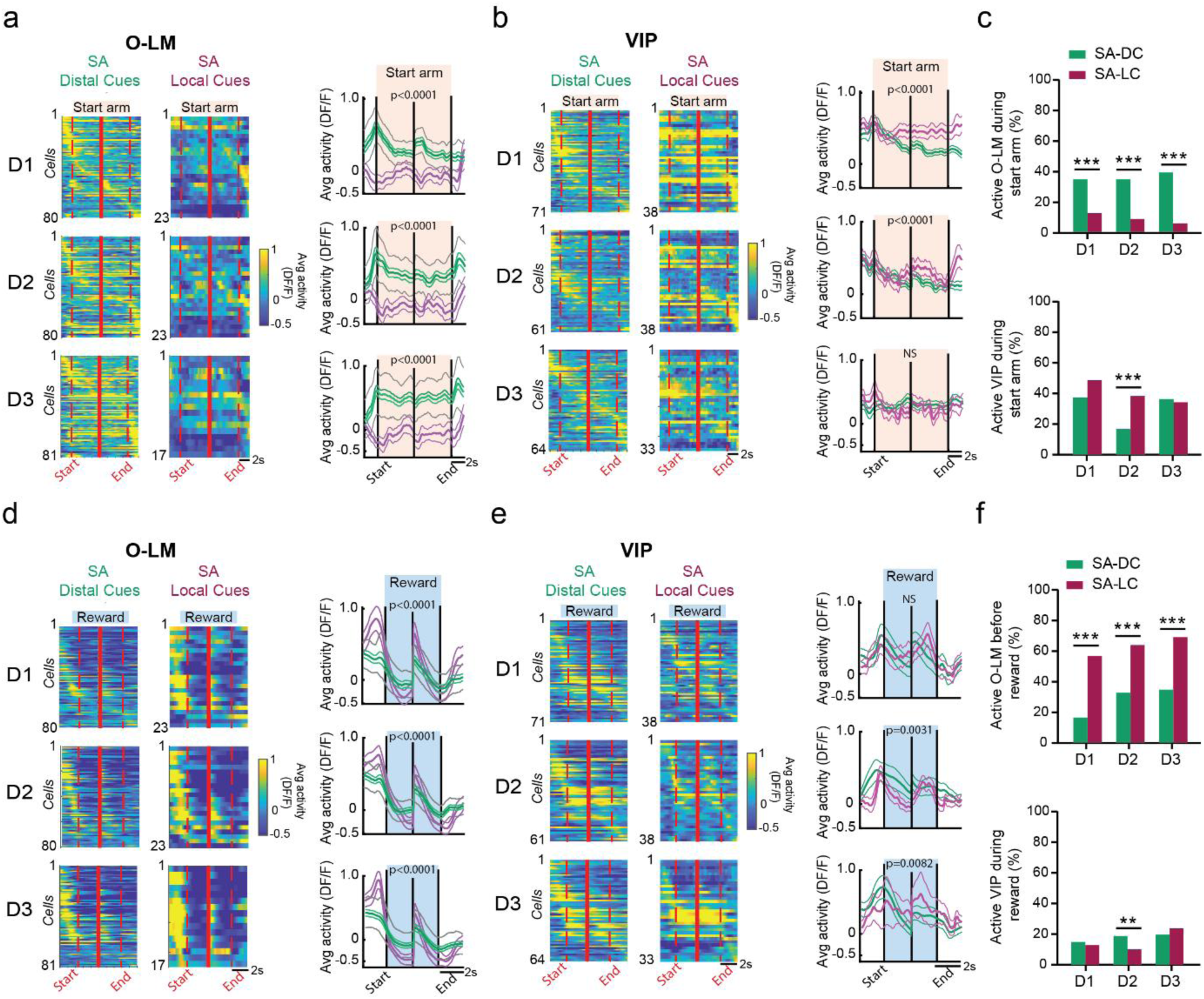
Distal cue availability in the start arm selectively modulates O-LM interneuron activity across learning but not VIP interneurons. **a,b,d,e**, Left: peri-event time histograms (PETHs) of averaged DF/F activity around the location phase **a-b,** or the reward consumption **d-e,** for both Star maze paradigms. Each heat map comprises two PETHs separated by a solid red line: one triggered by the start of the event (left dashed line) and one triggered by the end of the event (right dashed line). Each row represents the average activity of an individual cell. Right Panels: Mean (bold lines) and SEM (regular lines) population DF/F traces for each PETH for SA-DC (green) and SA-LC (magenta). The gray envelope around SA-DC traces in (**a)** and (**e)** denote the 99th percentile confidence interval determined by 1000 decimation iterations. **a,** O-LM interneurons are strongly active in the start arm in the SA-DC condition, but this activation is diminished when visual access to distal cues is removed (SA-LC condition. Condition * time interaction: Day 1, F(224,22624)=4.521, p<0.0001; Day 2, F(232,23200)=1.853, p<0.0001; Day 3, F(225,21375)=1.660, p<0.0001; α=0.05). **b,** VIP interneurons maintained (Day 3) or increased (Days 1 and 2) their activity during the start arm phase even without visual access to distal cues. Condition * time interaction: Day 1, F(299,31993)=3.316, p<0.0001; Day 2, F(295,28615)=2.829, p<0.0001; Day 3, F(283,26885)=2.010, p=0.2750; α=0.05). **c,** Top: The fraction of significantly active O-LM interneurons (determined by time-shuffling analysis) during the start arm was reduced in the SA-LC condition: (Day 1: 35% vs. 13%, x2=12.0888, p<0.0001; Day 2: 35% vs. 9%, x2=18.2110, p<0.0001; Day 3: 39% vs. 6%, x2=29.3620, p<0.0001). Bottom: The fraction of significantly active VIP (determined by time-shuffling analysis) during the start arm was unchanged (Day 1 and Day 3) or increased (Day 2) in the SA-LC condition (Day 1: 36% vs. 47%, x2=2.0595, p=0.1513; Day 2: 16% vs. 37%, x2=10.2683, p=0.0014; Day 3: 35% vs. 33%, x2=0.0223, p=0.8813). **d,** O-LM interneurons are inhibited during reward consumption in the SA-DC condition but are transiently activated just before mice reach the reward in the SA-LC condition. Condition * time interaction: Day 1, F(115,11615)=5.560, p<0.0001; Day 2, F(121,12100)=4.892, p<0.0001; Day 3, F(127,12065)=7.179, p<0.0001; α=0.05). **e,** VIP interneurons maintain their activity during reward consumption except for a slight reduction in the SA-LC condition. Condition * time interaction: Day 1, F(153,19737)=1.005, p=0.4664; Day 2, F(152,14744)=1.345, p=0.0031; Day 3, F(150,14250)=1.301, p=0.0082; α=0.05). **f,** Top: The fraction of significantly active O-LM interneurons before reaching the reward increased in the SA-LC condition: (Day 1: 16% vs. 56%, x2=33.0078, p<0.0001; Day 2: 32% vs. 63%, x2=18.0451, p<0.0001; Day 3: 34% vs. 69%, x2=23.1408, p<0.0001). Bottom: The fraction of significantly active VIP interneurons during reward consumption was unchanged in the SA-LC condition except for a reduction on Day 2 (Day 1: 36% vs. 47%, x2=2.0595, p=0.1513; Day 2: 16% vs. 37%, x2=10.2683, p=0.0014; Day 3: 35% vs. 33%, x2=0.0223, p=0.8813). **Statistics: a,b,d,e,** Two-way repeated-measures ANOVA. **c,f,** Chi2 proportion test with Yates correction.

To ensure that the smaller number of cells sampled in the SA-LC condition did not account for the differences reported, we performed a decimation analysis (see **Methods**) on the SA-DC datasets (see **Supplementary Fig. 3**). Briefly, we iteratively selected subsets of SA-DC cells to match the number of SA-LC cells and ran correlation analyses to compare these subsets with the original SA-DC and SA-LC data. This analysis, along with the 99% confidence intervals (gray bands, **Fig. 2a,d**), indicated that reduced sample size was not responsible for the observed differences. These findings demonstrate that the processing of distal cues during the start arm location phase strongly influences O-LM interneuron activity, while VIP interneurons were only marginally affected.

### Interneurons dynamics in relation to the navigation mode

To better understand how O-LM and VIP interneurons are modulated during the navigation phase (the interval between the trial start and reward), we analyzed the dynamic of their engagement across days. We hypothesized that interneurons involved in novelty detection would show higher firing rates during the initial session, as suggested by Tamboli et al.^59^. In contrast, if cells were more sensitive to spatial information processing, their activity would differ between the two Star maze conditions (SA-DC vs. SA-LC). We first examined changes in calcium transient rate (Hz) recorded during trials from both Star maze procedures (excluding the Start arm location phase). In the SA-DC condition, distinct dynamics emerged. VIP interneuron firing rates were significantly enhanced during Day 1, particularly in the early trials, but their activity was more stable by Day 2 and 3. In contrast, O-LM interneurons showed sustained activity throughout all sessions, even as VIP engagement waned (**Fig. 3a**, left and see **Supplementary Fig. 4a**). In the SA-LC paradigm, VIP interneuron transient frequency was increased on Day 1, whereas O-LM firing rates remained higher than in the SA-DC condition throughout all sessions. In the SA-LC condition, O-LM interneuron activity was the highest during the early phase of learning on Days 1 and 2. By Day 3, O-LM activity remained elevated throughout the entire session (**Fig. 3a**, right). This progressive increase in calcium event rate suggests that O-LM interneurons become increasingly engaged during navigation relying on a local frame of reference. VIP interneuron engagement in the SA-LC condition outlasted the first few minutes of testing on Day 1, remaining active throughout the day. This indicates that under complex conditions—where the experimental rules and context are difficult to extract— VIP interneurons remain engaged for an extended period, possibly to support a more complex familiarization with the experimental context (see **Supplementary Fig. 4a**).

**Fig. 3:**
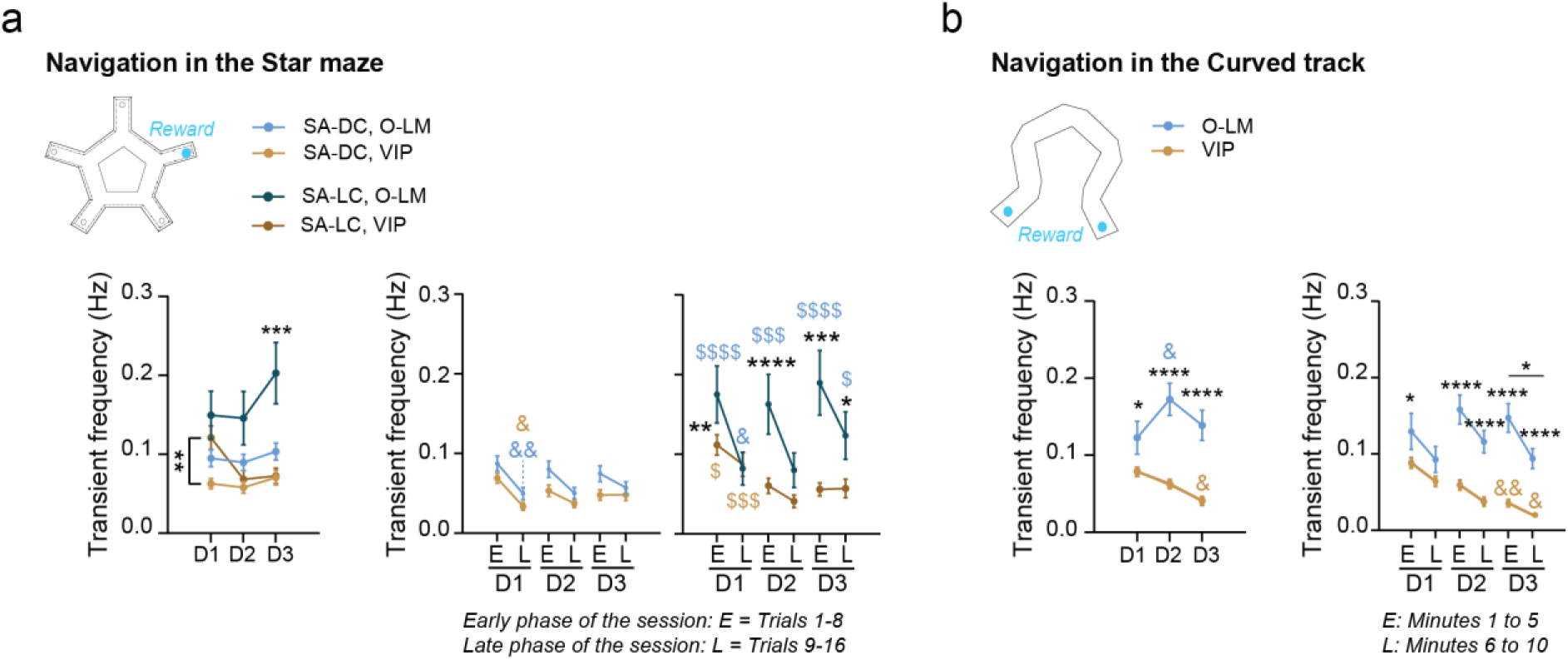
O-LM engagement depends on the navigation mode. **a,** Calcium transient frequency during the navigation phase of the Star maze procedures. Left: O-LM interneurons were more active during navigation than VIP interneurons (cell type effect: F(1,251)=41.60, p<0.0001; α=0.05). In the SA-LC condition, O-LM interneurons increased their activity level on Day 3 whereas VIP interneurons increased their activity on Day 1 (session * cell type* condition interaction F(2,251)=3.100, p=0.0468; α=0.05). Right panel: when accounting for intra-session learning dynamics (E: early and L: late), O-LM increased their activity in the SA-LC condition during early learning of each session, whereas VIP increased their activity only on Day 1 (phase * cell type * condition interaction F(2,251)=3.100, p=0.0468; α=0.05). **b,** Calcium transient frequency in the curved track. Left panel: O-LM were consistently more active during curved track navigation than VIP. VIP showed a progressive decrease in activity across days (cell type * session interaction: F(2,369)=4.750, p=0.0092; α=0.05). Right panel: when analyzing the first 5 minutes (early: E) vs. minute 6 to 10 (late: L) in the curved track, O-LM were more active across sessions during early training of each testing day, whereas VIP continuously decreased their activity levels (cell type * phase interaction: F(5,614)=4.241, p=0.0008; α=0.05). **Statistics: a-b,** 3-way ANOVA with LSD multiple comparisons post hoc test. Differences between conditions for the same genotype, same measure, same day $ p<0.05, $$ p<0.01, $$$ p<0.001 and $$$$ p<0.0001. Differences from Day1, for the same genotype, same measure: & p<0.05 and && p<0.01. Differences between groups on the same day: * p<0.05, ** p<0.01, *** p<0.001 and, **** p<0.0001. Data are shown as mean ± SEM.

To confirm that navigation based on a local frame of reference enhances O-LM interneurons firing rate during navigation, mice were tested in a linear “curved Track” with 16.3 cm high opaque walls that curtailed access to distant cues, compelling navigation through a local frame of reference (**Fig. 3b**). The curved shape design prevented mice from directly visualizing the reward port location, further favoring the use of a local reference frame. Mice freely explored the track for approximately 28-32 trials per day during three consecutive sessions. We found that O-LM interneurons remained dynamically engaged across all testing days, exhibiting firing rate levels similar to those recorded in the SA-LC version of the star maze (**Fig. 3b** **and see Supplemental Fig. 4b**). Consistent with the findings of Tamboli and collaborators^59^, VIP interneurons were particularly reactive during the first minute of training on Day 1, likely reflecting a response to novelty. Re-exposure to the curved track on Days 2 and 3 was accompanied by a progressive decrease in VIP firing rate, reaching a minimum on the last training session. This observation further supports the involvement of VIP interneurons in processing novelty, particularly when mice navigated based on a local frame of reference.

### O-LM and VIP place fields

Although spatial tuning is traditionally associated with principal cells^2^, several studies indicate that subsets of CA1 interneurons also exhibit place-cell properties^35,60–63^. However, whether this spatial tuning is modified by the navigational or cognitive strategies an animal employs has not been determined. To address this, we performed place field analyses on O-LM and VIP interneurons in both the curved track and Star maze protocols. We applied a conservative split-half stability shuffling analysis to identify place cells with stable spatial firing^64^ (see **Methods**). In the curved track, where local cues were prominent, about 20% of O-LM interneurons and 10% of VIP interneurons met place cell criteria (**Fig. 4a,b**). O-LM place fields were primarily located in the maze alley, where mice actively navigate while VIP place fields clustered near the reward region (**Fig. 4c**). In the Star maze, when distal cues were available (SA-DC), around 10% of O-LM and VIP interneurons qualified as place cells (**Fig. 4d,e**), with O-LM place fields predominantly localizing close to decision-making points (**Fig. 4f**). However, under conditions favoring a local reference frame (SA-LC), O-LM interneurons showed a higher proportion of place cells on Day 1 (approximately 25%) but their place fields location was shifted toward the maze’s inner alley. The chosen frame of reference also influenced VIP place fields, shifting them from target arms to non-target (error) arms, mirroring the animals’ difficulty in locating the target arm (**Fig. 4d-f**). These dynamic shifts underscore the remarkable adaptability of interneurons in modifying their spatial encoding properties under the control of the reference frame.

**Fig. 4:**
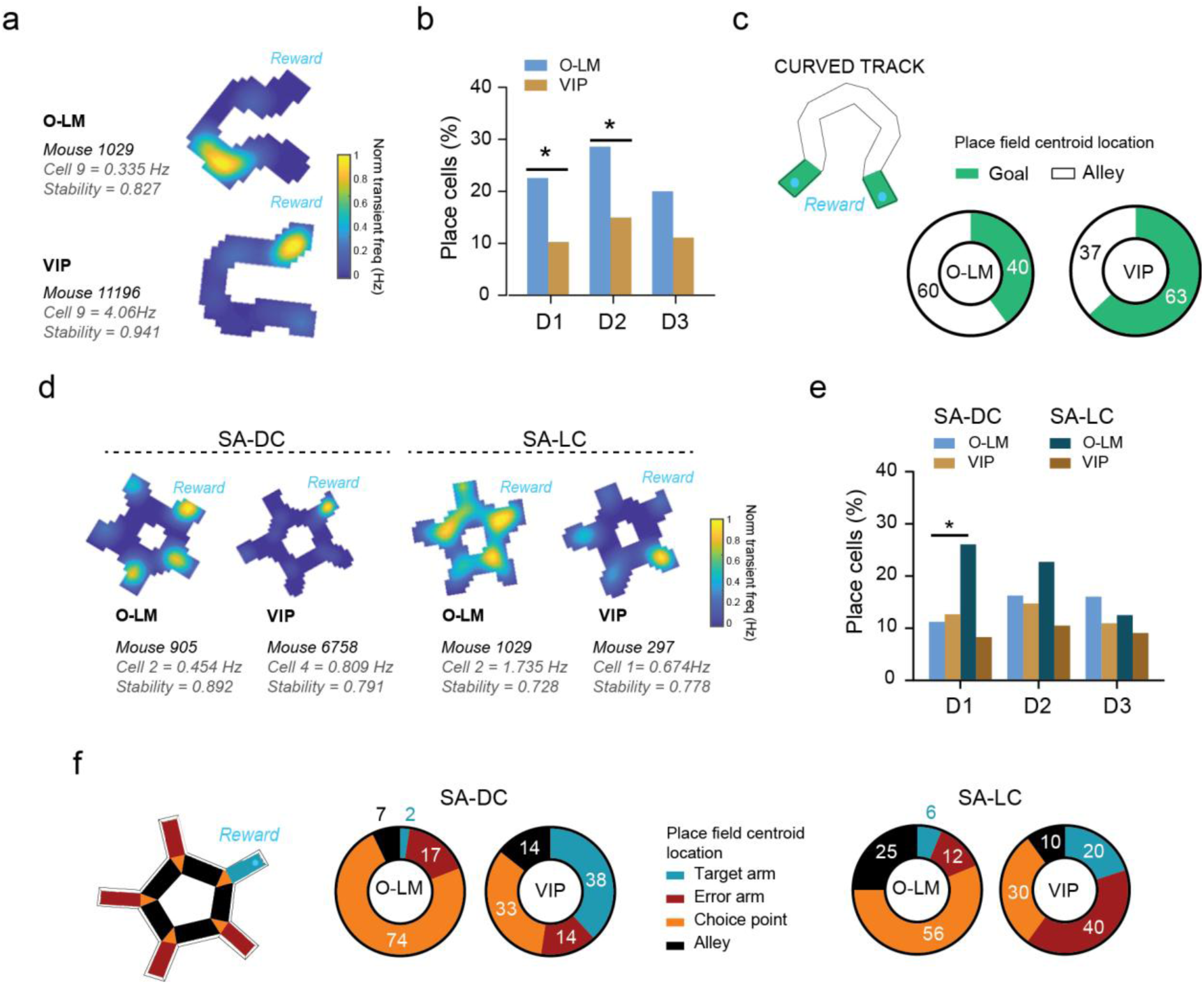
Influence of the frame of reference on O-LM and VIP place field location. **a,** Representative place cells recorded from the curved track, with their within-field peak activity and stability measure. **b,** In the curved track, the fraction of interneurons classified as place cells was greater in O-LM than in VIP cells, especially on the first two sessions (Day 1: 23% vs. 10%, x2 = 5.2224, p=0.0223; Day 2: 29% vs. 15%, x2 = 4.9242, p=0.0265; Day 3: 20% vs. 11%, x2 = 2.4432, p=0.1180). **c,** Place cell field centroids were preferentially located in the alleys of the curved track for O-LM cells and in goal regions for VIP interneurons (60% vs. 37%, x2 = 9.6887, p=0.0019).**d,** Representatives place cells recorded from the Star maze SA-DC and SA-LC conditions. **e,** In the Star maze, the fraction of interneurons classified as place cells was similar between O-LM and VIP interneurons, except for an increased proportion of O-LM classified as place cells on Day 1 of the SA-LC condition (Day 1, O-LM place fields: SA-DC versus SA-LC: 26% vs. 11%, x2 = 6.4998, p=0.0108). **f,** In the Star maze, place fields centroids were mostly located in choice points for O-LM interneurons and in arms for VIP cells. Removal on distal cues in the start arm affected both cell types: O-LM proportion of place fields centroids located in the inner alley of the maze increased (25% vs. 7%, x2 = 10.715, p=0.001), VIP proportion of place fields centroid located in non-target arms increased (40% vs. 14%, x2 = 15.855, p<0.0001). **Statistics: b,e,** 3-way ANOVA with Bonferroni multiple comparisons post hoc test. Difference between condition, same session: * p<0.05. Data are shown as mean ± SEM.

Additional place field metrics are shown in **Supplementary Fig. 5**. Taken together, these findings demonstrate a strong influence of the frame of reference on both the number and location of place fields, reflecting O-LM place fields preference for regions linked to active navigation and VIP place fields preference for reward locations or potential reward zones.

### O-LM locomotion tuning as a function of the frame of reference

O-LM and VIP interneurons have been shown to increase their activity with locomotion^35,65^. However, it’s also been shown that a small proportion of interneurons are activated during immobility^32^. We hypothesize that the frame of reference plays a critical role in shaping the correlation between O-LM calcium activity and behavioral states, whereas VIP activity may remain stable across the different navigation modes. To determine this, we analyzed interneuron activity during the navigation phase of the Star maze by comparing immobility and locomotion states. Locomotion events were defined as periods of movement with a running speed over 5 cm/sec and lasting for at least 1 second while immobility events were defined as periods below this speed threshold (see **Methods**). Under SA-LC conditions, where navigation relies on a local frame of reference, the proportion of time spent in locomotion *versus* immobility progressively converged across training days (**Fig. 5a**). Notably, O-LM firing rates during locomotion increased across days, while VIP firing rates were elevated only on Day 1 (see **Supplementary Fig. 6c,d**). To quantify cell-state preference, we classified cells as preferentially modulated by immobility, locomotion, or neither (similar activity levels during locomotion and immobility) using a modulation score and a shuffling analysis (see **Methods**). In the SA-DC conditions (distal reference frame), similar proportions of O-LM cells were tuned to either locomotion or immobility (approximately 20% each) across all training days. VIP interneurons displayed a similar pattern on Day 1 but showed a marked reduction in immobility-preferring cells by Days 2 and 3 (**Fig. 5b,c**). Overall, when navigation depended on distal cues, both O-LM and VIP contained subsets of cells tuned to either locomotion or immobility. Remarkably, under SA-LC conditions, O-LM interneurons became solely locomotion-tuned, with an increased proportion as training progressed, emphasizing their strong engagement during active navigation (**Fig. 5b,c**). In contrast, VIP interneurons maintained consistent proportions of locomotion-tuned cells with only a few immobility-preferring cells on Days 1 and 3. These differences are not due to the difference in cell number between conditions as a decimation analysis on SA-DC datasets indicated that reducing the sampling size of the SA-DC condition to match SA-LC cell numbers did not yield similar proportions of locomotion-preferring O-LM cells as observed in this latter condition (see **Supplementary Fig. 7**).

**Fig. 5:**
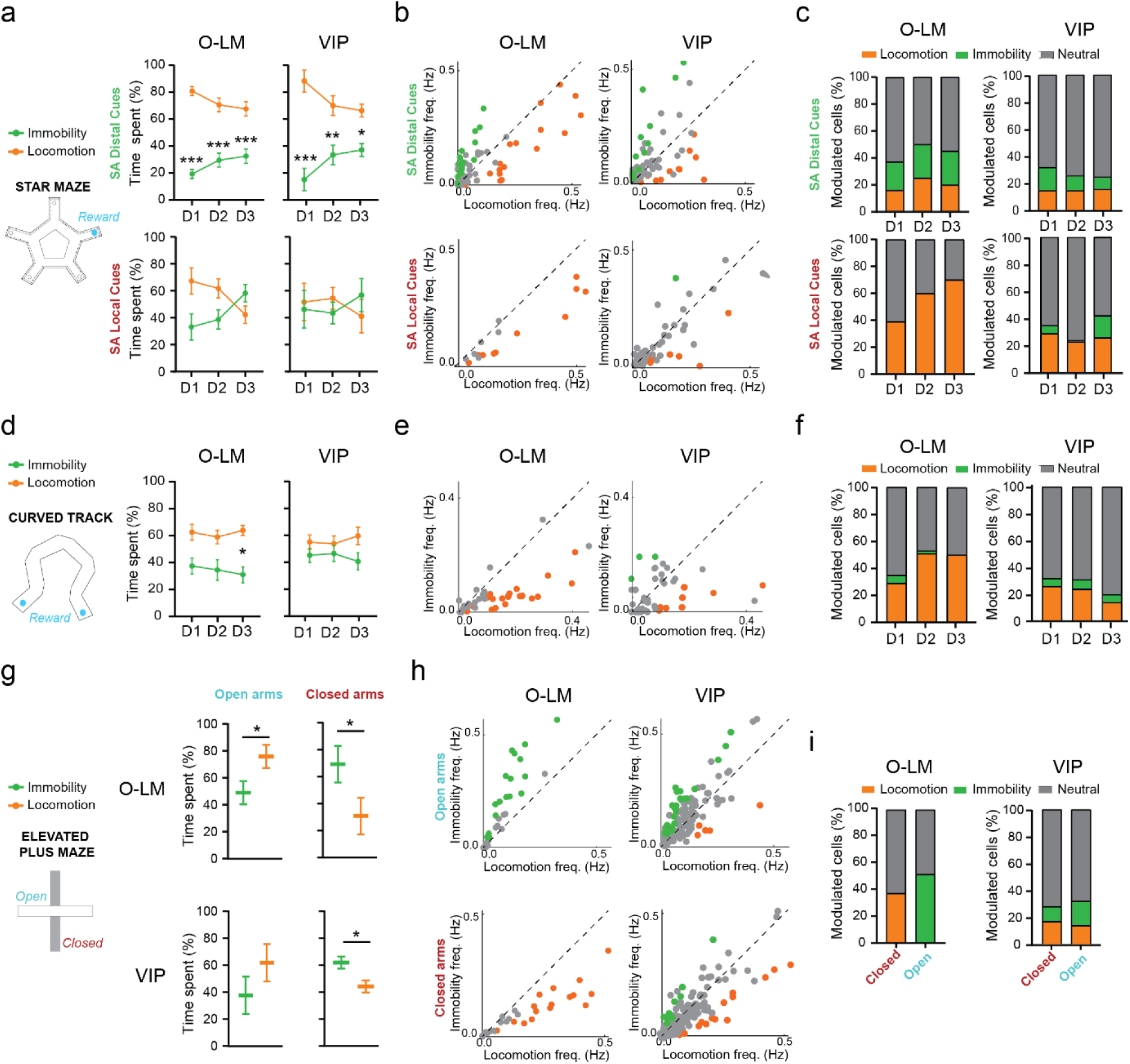
Bi-modal O-LM engagement during locomotion and immobility as a function of the frame of reference used during navigation. **a,** Left: experimental apparatus. Right: the proportion of time spent in locomotion and immobility during the navigation phase of the Star maze was affected by the condition for Chrna2 mice (condition * locomotion state interaction: F(1,18)=17.28, p=0.0006; α=0.05) and time (day * locomotion state interaction: F(2,27)=8.233, p=0.0006; α=0.05) but not by their interaction (day * locomotion state * condition interaction: F(2,18)=1.624, p=0.2246; α=0.05). Right: for VIP animals, the proportion of time spent in locomotion and immobility during the navigation phase of the Star maze was affected by the condition (condition * locomotion state interaction: F(1,12)=27.29, p=0.0002; α=0.05). **b,** Scatterplots of calcium activity during immobility and locomotion on Day 3. Each dot represents an individual interneuron, dots are color-coded according to cell classification: significantly preferring locomotion (orange), immobility (green) or with no significant preference (gray), as determined by time-shuffling analysis. **c,** Fractions of O-LM and VIP interneurons. Left: the proportion of O-LM neurons that prefer locomotion increased across days in the SA-LC condition (Day 1: 39% vs. 16%, x2 = 12.1379, p=0.0004; Day 2: 60% vs. 25%, x2 = 23.6522, p<0.0001; Day 3: 70% vs. 20%, x2 = 48.5051, p<0.0001). Right: the proportion of VIP interneurons that prefer locomotion increased in SA-LC only on Day 1 (Day1: 29% vs. 15%, x2 = 4.9242, p=0.0265; Day 2: 23% vs. 15%, x2 = 1.5919, p=0.207; Day 3: 26% vs. 16%, x2 = 2.4412, p=0.1182). **d,** The proportion of time spent in locomotion and immobility was not different across genotypes in the curved track (cell type * locomotion state interaction: F(1,50)=2.789, p=0.1012; cell type * locomotion state * session interaction: F(2,50)=0.06579, p=0.9364; α=0.05). **e-f,** Scatterplots for Day 3 (**e)** and corresponding fractions of O-LM and VIP cells (**f)** significantly modulated by locomotion (orange), immobility (green) or with no significant preference (gray) as determined by time-shuffling analysis. **e,** Each dot represents an individual interneuron. **f,** O-LM preference for locomotion was higher than VIP starting from Day 2 (Day 1: 29% vs. 26%, x2 = 0.1003, p=0.7515; Day 2: 51% vs. 24%, x2 = 14.4213, p=0.0001; Day 3: 50% vs. 14%, x2 = 28.1480, p<0.0001). **g,** In the EPM, the proportion of time spent in locomotion and immobility was influenced by the type of arm in a similar way for both mice strain (arm * locomotion state interaction: F(1,16) = 74.34, p<0.0001; genotype * locomotion state * arm interaction: F(1,16)=1.509, p=0.2371; α=0.05). **h-i,** Scatterplot for EPM (**h)** and corresponding fractions of O-LM and VIP cells (**i)** significantly modulated by locomotion (orange), immobility (green) or with no significant preference (gray) as determined by time-shuffling analysis. **h,** Each dot represents an individual interneuron. **i,** O-LM preference shift from locomotion in closed arms to immobility in open arms, whereas VIP interneurons maintained similar proportion across arms (locomotion: 17% vs. 14%, x2 = 0.1527, p=0.6960; Immobility: 11% vs. 18%, x2 = 1.4519, p=0.2282). **Statistics: a,d,g,** 3-way ANOVA with Bonferroni multiple comparisons post hoc test. Difference between locomotion and immobility: * p<0.05, ** p<0.01, *** p<0.001. Data are shown as mean ± SEM.

To confirm that O-LM interneuron activation during locomotion is linked to a local-frame based navigation, we performed the same analysis in the curved linear track, which restricts distal cue use by tall walls (**Fig. 5d**). Consistent to the SA-LC results, O-LM interneurons in the curved track were preferentially and increasingly engaged during locomotion (Day 1: 29%; Day 2: 51%, and Day 3: 50%; **Fig. 5e,f**) while immobility-preferring O-LM cells progressively declined. VIP cell modulation resembled that observed in the SA-LC Star maze condition, but their firing rates during locomotion declined with re-exposure between Day 1 and subsequent sessions, suggesting a reduced engagement as the environment became more familiar. Therefore, O-LM interneurons exhibit heightened sensitivity to the frame of reference whereas VIP cells become less engaged as the task or environment becomes familiar.

To further test how immediate distal cue availability affects interneurons activity, we analysed the five minutes of spontaneous exploration in an Elevated Plus Maze (EPM). While this paradigm is traditionally used to assess anxiety, the EPM also provides distinct reference frame contexts: closed arms dominated by a local reference frame, and open arms allowing for distal reference frame utilization^66,67^. Estimation of the proportions of cells modulated by the animal’s behavioral states revealed that O-LM interneurons were preferentially tuned by locomotion when mice were in closed arms and by immobility when mice visited open arms (**Fig. 5h,i**). VIP interneurons, in contrast, exhibited stable proportions of locomotion- and immobility-tuned cells across both closed and open arms, indicating a lack of sensitivity to changes in the reference frame. A down-sampling control analysis confirmed that these effects were not attributable to differences in time spent in both arm types (see **Methods** and **Supplementary Fig. 8**). Overall, these results confirm that O-LM activity depends on the reference frame and distal cue availability, while VIP interneurons decrease their engagement as the environment becomes more familiar.

### O-LM bi-modal speed coding

Speed coding in the hippocampus is widely considered as a fundamental feature of interneuron function, characterized by stability across space, time, and context^25,35^. Most reports indicate that O-LM and VIP interneurons modulate their activity according to running velocity^32,35,65,68^. However, a recent study has suggested that O-LM interneurons may exhibit a negative correlation with speed^44^. This discrepancy led us to hypothesize that contextual factors, specifically the spatial frame of reference, may influence O-LM speed coding, whereas VIP speed coding would remain unaffected. We first confirmed that running speed was stable across sessions, genotypes, and experimental conditions (see **Supplementary Fig. 10**). We then analyzed how the interneuron firing rates relate to running speed (see **Methods**). Under the Star maze SA-LC conditions (local frame of reference), O-LM activity exhibited a robust positive correlation with running speed, with firing rates increasing proportionally to the speed increase (**Fig. 6a,b**). Conversely, under SA-DC conditions (distal frame of reference), O-LM activity showed a weaker or even negative correlation with speed, and average firing rates decreased with increasing running speeds. In contrast, VIP speed coding remained unchanged between conditions (**Fig. 6d,e**). To further investigate speed coding mechanisms, we performed a population vector (PV) analysis, pooling all cells into pseudo-populations (**Fig. 6c,f**). This analysis suggested that the speed coding of O-LM interneurons was bimodal: a population coding of the speed in the SA-DC condition relying on a distal frame of reference, and a rate-based coding of the speed in the SA-LC condition associated with a local frame of reference (**Fig. 6c**). In contrast, VIP cells displayed a population coding of the speed in both conditions (**Fig. 6f**).

**Fig. 6:**
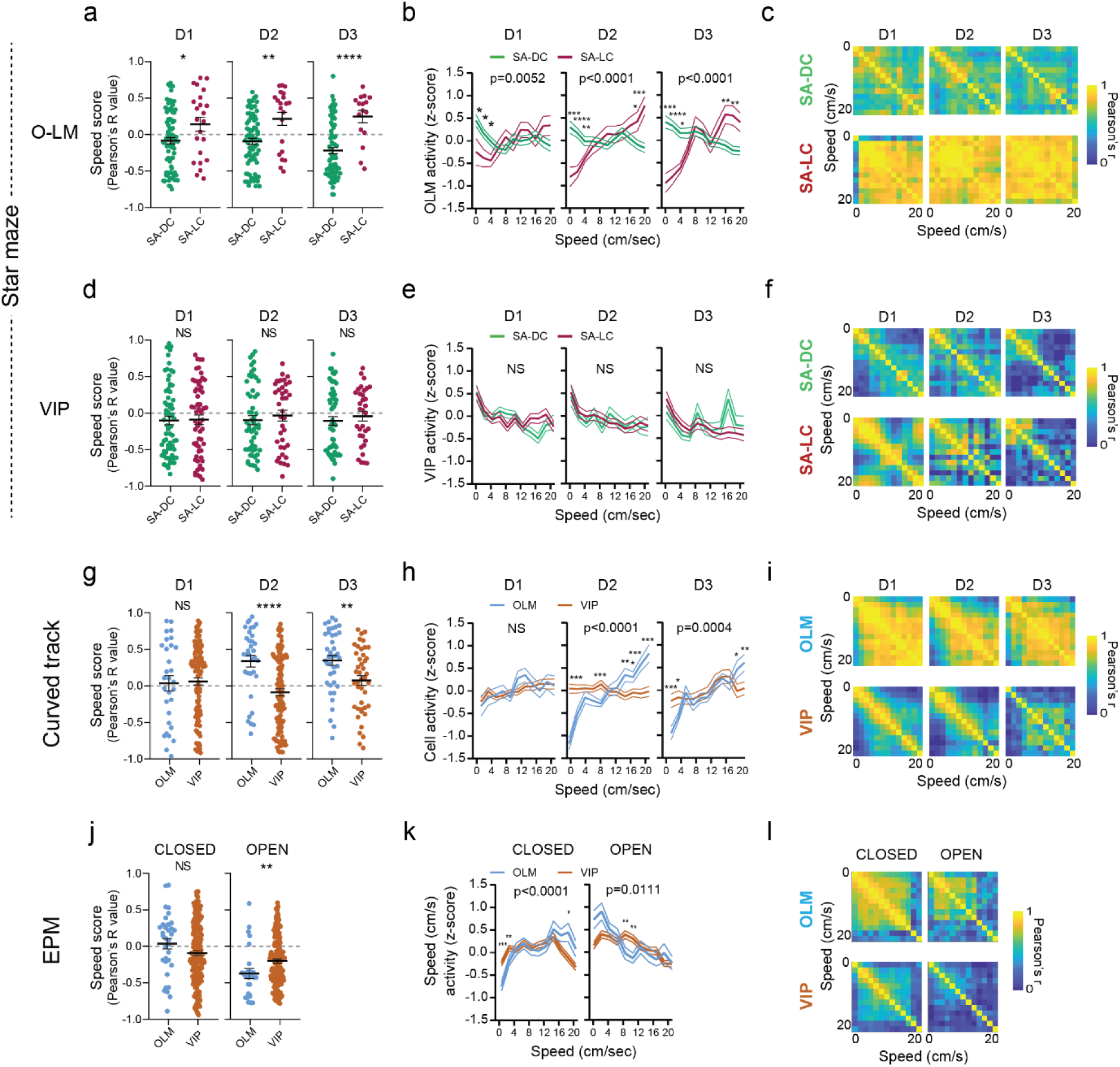
Bi-modal O-LM speed coding properties depending on the frame of reference. **a,** In the Star maze, O-LM speed scores were higher in the SA-LC condition (Day 1: t(101)=2.298, p=0.0236 ; Day 2: t(97)=3.294, p=0.0014 ; Day 3: t(90)=4.471, p<0.0001; α=0.05). **b,** O-LM average activity decreased with increasing running speed in the SA-DC condition whereas it ramp up with increasing speed in the SA-LC condition (condition * speed interaction: Day 1: F(12,1212)=2.368, p=0.0052 ; Day 2: F(12,1176)=5.520, p<0.0001 ; Day 3: F(12,1092)=7.542, p<0.0001; α=0.05). **c,** O-LM cells pseudo population vectors (PV) suggest a more orthogonal representation of the speed in the SA-DC condition than in the SA-LC condition, suggesting a population code of the speed in the SA-DC condition and a rate-coding of the speed in the SA-LC condition. **d,** In the Star maze, VIP speed scores did not differ between condition (Day 1: t(129)=0.1187, p=0.9057 ; Day 2: t(93)=0.6388, p=0.52445 ; Day 3: t(88)=0.6884, p=0.4930; α=0.05). **e,** VIP average activity tends to decrease with speed similarly in both conditions (condition * speed interaction: D1: F(10,1290)=1.769, p=0.0616 ; D2: F(10,970)=0.4294, p=0.9327 ; D3: F(10,950)=1.047, p=0.4014; α=0.05). **f,** VIP interneuron PV suggests a similar orthogonal representation of the speed consistent with a population coding of the running speed. **g,** In the curved track, where proximal cues are prominent, O-LM cells speed scores were higher than VIP starting from Day 2 (Day 1: t(144)=0.2207, p=0.8256 ; Day 2: t(138)=4.341, p<0.0001 ; Day 3: t(81)=2.854, p=0.0055; α=0.05). **h,** In contrast to VIP cells, O-LM interneurons increased their activity with increasing speed starting from Day 2 in the curved track (cell type * speed interaction: D1: F(10,1430)=0.6202, p=0.7977 ; D2: F(10,1380)=9.603, p<0.0001 ; D3: F(10,810)=3.249, p=0.0004; α=0.05). **i,** PV analysis suggests that in the curved track O-LM rate-codes the running speed whereas VIP interneurons display a population coding of the running speed. **j,** In the EPM, O-LM speed scores were lower than VIP in open arms but similar in closed arms (open: t(188)=2.411, p=0.0169 ; closed: t(200)=1.653, p=0.0999 ; α=0.05). **k,** In contrast to VIP, O-LM interneurons respond differentially to speed in each arm type, increasing their activity with speed in closed arms whereas decreasing their activity with increasing speed in open arms (speed * arm * cell type interaction: F(10,2035) = 4.150, p<0.0001; α=0.05) **l,** PV analysis suggests that VIP maintained a population coding of the speed in both arms whereas O-LM cells displayed a rate coding of the speed in the closed arm. **Statistics: a,d,g,j,** two-tailed unpaired t test: * p<0.05, ** p<0.01, *** p<0.001, **** p<0.0001. **b,e,h,** 2-way ANOVA with Bonferroni multiple comparisons post hoc test. **k,** 3-way ANOVA with Bonferroni multiple comparisons post hoc test. Difference between SM conditions or cell type: * p<0.05, ** p<0.01, *** p<0.001. Data are shown as mean ± SEM.

To validate that O-LM rate-coding of the speed is characteristic of a local frame of reference, we analyzed data from the curved track, which restricts distal cue use (**Fig. 6g-i**). Similar to the SA-LC condition, O-LM activity on the curved track was increasingly positively correlated with speed starting from Day 2 (**Fig. 6g**), and average firing rates ramped up with increasing speed (**Fig. 6h**). The PV analysis of O-LM cells also indicated a rate-coding of the running speed in this local frame of reference (**Fig. 6i**). VIP neurons showed similar speed coding patterns to those observed in the Star maze, maintaining population coding regardless of frame of reference.

To further explore the influence of the frame of reference on O-LM interneurons, we analyzed their activity in the Elevated Plus Maze (EPM). A higher correlation between O-LM activity and speed was found in closed arms compared to open arms (**Fig. 6j**). In closed arms, O-LM firing rates ramped up with increasing speed, while in open arms (where distal cues were available), activity decreased at higher speeds (**Fig. 6k**). PV analysis supported again a rate-based code under local frames of reference (closed-arm) and a population-based code under distal frame of reference (open arms, **Fig. 6l**). VIP interneurons maintained stable speed coding across both arm types, unaffected by the frame of reference. Down-sampling the closed-arm data to match open-arm visits duration confirmed that those differences in O-LM speed coding properties depend on the frame of reference and were not caused by shorter visits in the open arms (see **Supplementary Fig. 9**).

Altogether, these results suggest that O-LM interneurons exhibit a bi-modal speed-coding mode: under the distal frame of reference (Star-maze SA-DC, EPM open arms), they display a population-based mode where activity peaks at lower speeds, whereas under local frames of reference, their firing rates scale up with increasing speed in a rate-coding mode. In contrast, VIP interneurons exhibit a more canonical context-independent, population-based speed coding strategy, maintaining stable encoding of the speed across all conditions.

### Navigation strategy and contextual information coding

Beyond speed and spatial coding, interneurons may also integrate behavioral information about the task structure, especially in complex cognitive tasks such as reference memory in the Star maze. To investigate this hypothesis, we measured the average O-LM and VIP interneuron activity across the main phases of the task: the four start arm locations relative to the target arm position (−2, -1, +1, +2), the different behavioral states during the trial (immobility, locomotion in the center of the maze, trajectories to error arms, trajectories to the target arm), reward consumption, and post-reward behaviors (immobility and locomotion, see **Method** and **Fig. 7a,b**). In the SA-DC condition, O-LM interneurons activity peaked during the critical location phase in the start arms, whereas in the SA-LC condition, O-LM activity shifted predominantly to the trial phase, with maximal activation during locomotion (**Fig. 7a,c**). This was reproducible during the three learning days. Such a difference was not observed for VIP interneurons (**Fig. 7b,d**). We then constructed pseudo-population vectors of the different behavioral phases to analyze the orthogonality of O-LM and VIP population activity (see **Supplementary Fig. 11**). Across the three learning days, O-LM PV tended to become more orthogonal in both conditions, but the degree of population activation similarity across the different behavioral phases was higher for the SA-LC condition, suggesting more overlap between the activity during the different behavioral phases in this condition. On the other hand, VIP interneurons maintained a similar level of orthogonality across conditions, except for more similar activity for immobility and locomotion events that did not lead to the target arm. Thus, successful navigation based on a distal frame of reference in the SA-DC condition was associated with an increasing behavior-specific activation of O-LM at the population level, whereas navigation based on a local frame of reference dramatically reduced this behavior-specific modulation, in favor of locomotion and speed rate-coding. Taken together, this additional functional divergence underscores the specialized roles of O-LM and VIP interneurons in integrating task structure and navigation strategies.

**Fig. 7:**
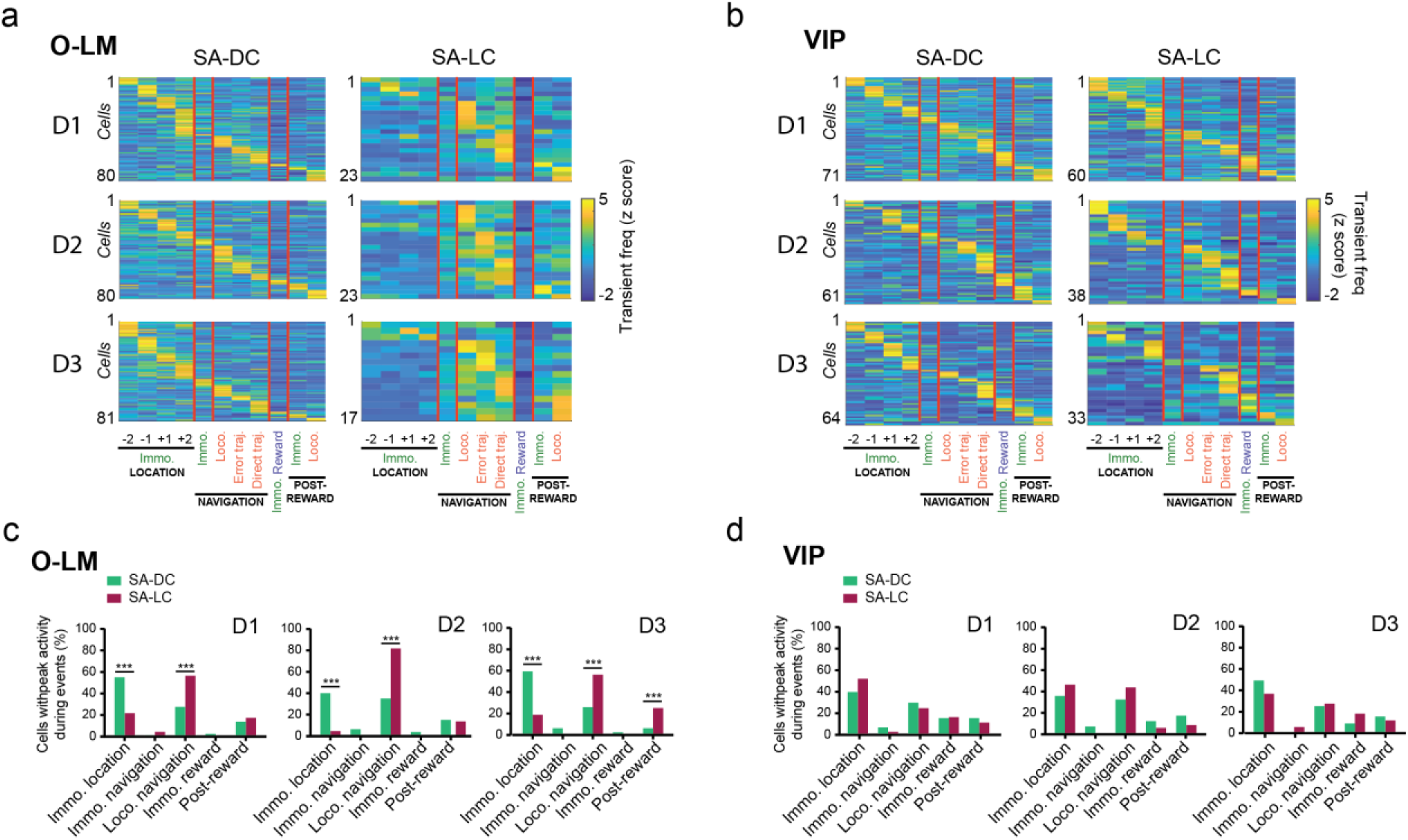
O-LM activation across behavioral events depends on the frame of reference. **a-b,** Z-scored average transient frequencies of O-LM (**a**) and VIP (**b**) interneurons in the SA-DC (left) and SA-LC conditions (right). Analysis was performed for immobility periods (green) during the location periods into the -2, -1, +1, and +2 start arms as well as during reward consumption and post-reward period. Analysis for locomotion periods (orange) was made during navigation (ending in the center of the maze), during error trajectories (locomotion ending in an error arm), during direct trajectories (locomotion ending in the target arm), and during post-reward. **a,** O-LM peak activity spreads across all behavioral events in the SA-DC condition but concentrates on locomotion in the SA-LC condition. **b,** VIP interneuron peak activity was evenly distributed across all behavioral events in both Star maze conditions. **c,** Fractions of O-LM cells having their activity peak during the different phases of the task for both conditions. The SA-DC condition was associated with a higher fraction of OLM interneurons preferentially active during the start arm location phase (Day 1: 55% vs. 22%, x2 = 21.6239, p<0.0001; Day 2: 40% vs. 5%; x2 = 33.147, p<0.0001; Day 3: 59% vs. 19%; x2 = 31.9672, p<0.0001) whereas the SA-LC condition was associated with higher OLM interneuron activation during locomotion (Day 1: 28% vs. 56%, x2 = 14.9631, p=0.00011; Day 2: 35% vs. 81%, x2 = 41.564, p<0.0001; Day 3: 26% vs. 56%, x2 = 17.383, p<0.0001). **d,** The fractions of VIP interneurons that had their peak of activity during the different phases of the task were not different in the SA-LC vs. SA-DC conditions. **Statistics:** Chi2 proportion test with Yates correction.

### O-LM speed tuning allows decoding of the navigational frame of reference

To explore how O-LM and VIP interneuron activity patterns contained information about the dominant reference frame used by mice, we performed a linear decoding analysis using a support vector machine (SVM). Speed scores were selected as the decoding variable because of their sensitivity to the reference frame (**Fig. 6**). Given that each mouse contributed relatively few cells, and their numbers varied across sessions, we combined all cells into pseudo-populations and used the daily average speed scores as the SVM input variable (see **Methods**). Sessions were labeled as relying on either proximal reference frame (EPM closed arm, curved track, SA-LC condition of the star maze) or a distal reference frame (EPM open arm, SA-DC condition of the star maze). To avoid overfitting, we excluded the test session to decode from its corresponding training dataset. Using this approach, we found that O-LM but not VIP speed scores provided a reliable and accurate index of the dominant reference frame used by mice (**Fig. 8**). O-LM decoding performance improved with training (**Fig. 8c-e**), suggesting that in a new environment, mice may initially use a mixture of both reference frames. Neither O-LM nor VIP speed scores could decode the reference frame in the EPM closed arms, although both successfully classified the open arms (**Fig. 8b**). This may be due to the size of the maze: a mouse in the closed arm can still get a decent visual access to distal cues through the center choice point formed by the intersection of closed and open arms.

**Fig. 8:**
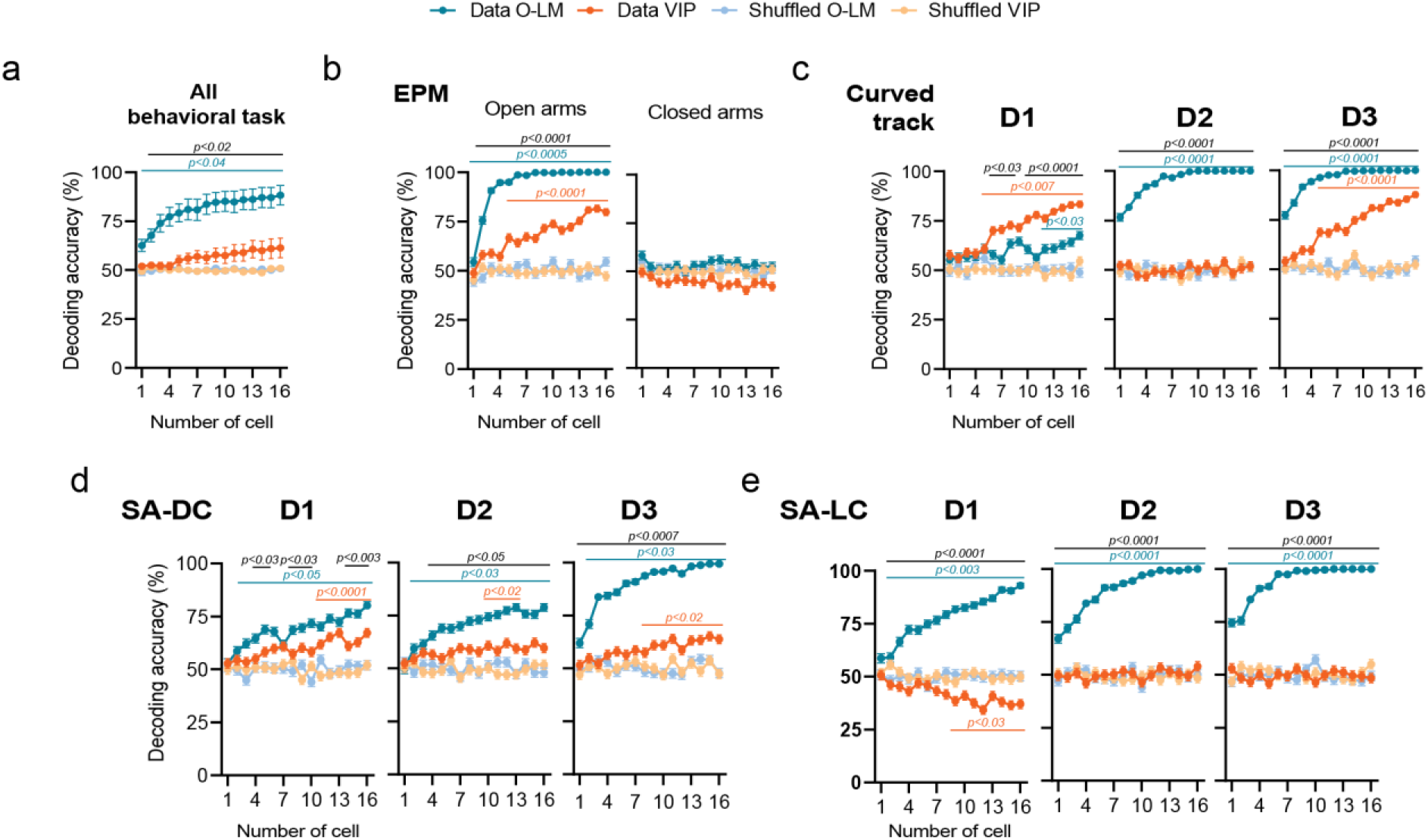
O-LM speed scores, unlike VIP, reliably decode the reference frame across different tasks. **a**, When data from all behavioral tasks were combined, decoding the reference frame using interneuron speed scores was more accurate for O-LM than VIP (cell type × cell number interaction: F(15,300) = 5.355, p < 0.0001). **b,** Left: In the EPM open arm, both O-LM and VIP speed scores decoded the reference frame as distal, though O-LM decoding was more accurate (cell type × cell number interaction: F(15,15968) = 13.00, p < 0.0001). Right: In the closed arm, neither O-LM nor VIP speed scores could decode the reference frame as proximal, regardless of the number of cells (cell type × cell number interaction: F(15,15968) = 0.5835, p = 0.8900). **c,** Left (Day 1 on the curved track): Both O-LM and VIP speed scores decoded the reference frame as proximal, though VIP showed lower accuracy (cell type × cell number interaction: F(15,15968) = 6.823, p < 0.0001). Middle (Day 2): Only O-LM speed scores decoded the frame of reference as proximal (cell type × cell number interaction: F(15,15968) = 10.26, p < 0.0001). Right (Day 3): Both O-LM and VIP speed scores decoded the reference frame as proximal, but O-LM-based decoding was more accurate (cell type × cell number interaction: F(15,15968) = 10.52, p < 0.0001). **d,** In the SA-DC condition of the star maze, both O-LM and VIP speed scores decoded the reference frame as distal, although O-LM-based decoding was consistently more accurate across all three sessions: Day 1 (F(15,15968) = 2.186, p = 0.0051), Day 2 (F(15,15968) = 3.968, p < 0.0001), and Day 3 (F(15,15968) = 8.631, p < 0.0001). **e,** In the SA proximal cue condition, only O-LM speed scores successfully decoded the reference frame as proximal. VIP speed scores incorrectly classified it as distal on Day 1 (F(15,15968) = 27.34, p < 0.0001), Day 2 (F(15,15968) = 15.62, p < 0.0001), and Day 3 (F(15,15968) = 13.19, p < 0.0001). **Statistics:** Data were analyzed using two-way ANOVA, followed by Sidak’s multiple comparisons. Significant differences between O-LM and VIP data are shown in black, between O-LM and shuffled O-LM data in blue, and between VIP and shuffled VIP data in orange.

### O-LM interneurons support spatial reference memory formation

To investigate whether O-LM interneurons contribute to efficient spatial reference memory, we focused on the SA-DC paradigm, where O-LM cells displayed a reliable activation at the starting point and a strong modulation during immobility. Moreover, a subset of these interneurons formed place fields at critical maze intersections, particularly at choice points. Based on these observations, we examined the impact of silencing O-LM interneurons on the full Star maze procedure. We used an inhibitory DREADD receptor (AAV2-hSyn-DIO-hM4D(Gi)-mCherry) or a control virus (AAV2-Ef1α-flex-tdTomato) targeted to O-LM interneurons (see **Methods**). Post-hoc histology confirmed a strong DREADD receptor expression in the stratum lacunosum moleculare, where O-LM axons ramify (**Fig. 9a**). To activate DREADD receptors, mice consumed 2 mg/kg of Compound 21 (C21) diluted in sucrose water, a dose selected based on pharmacokinetic and functional analyses (**Fig. 9b,c**). Calcium imaging revealed a significant, albeit incomplete, reduction in O-LM activity two hours after C21 ingestion. We then assessed locomotion, exploration, and spatial reference memory (**Fig. 9d-f**). Mice expressing DREADD-hM4Di performed less accurately on Days 1 and 2 of the SA-DC Star Maze version, committing more errors in locating the target arm, particularly during the first eight trials. By Day 3, their performance matched that of controls, indicating delayed, rather than impaired, learning. Neither the navigational strategy (**Fig. 9e**) nor speed (see **Supplementary Fig. 12**) was affected by O-LM silencing. During memory retrieval testing, conducted without C21 administration, both groups accurately located the rewarded arm (**Fig. 9f**). Thus, despite their relatively small population, dorsal O-LM interneurons were essential for an optimized spatial reference learning, but their inactivation did not disrupt memory retrieval once the task was mastered. Next, we examined the effect of O-LM inhibition on local cue-based navigation. The same mice were trained on the curved Track and in the EPM (see **Supplementary Fig. 11**). Across training days, chemogenetic inhibition of O-LM cells did not substantially affect locomotion states. During free exploration of the EPM, DREADD-hM4Di-expressing mice showed no differences in open arm visits, speed, or total locomotion time. These findings indicate that silencing O-LM interneurons does not affect anxiety or locomotion performance but rather delays spatial learning under distal cue–based conditions.

**Fig. 9:**
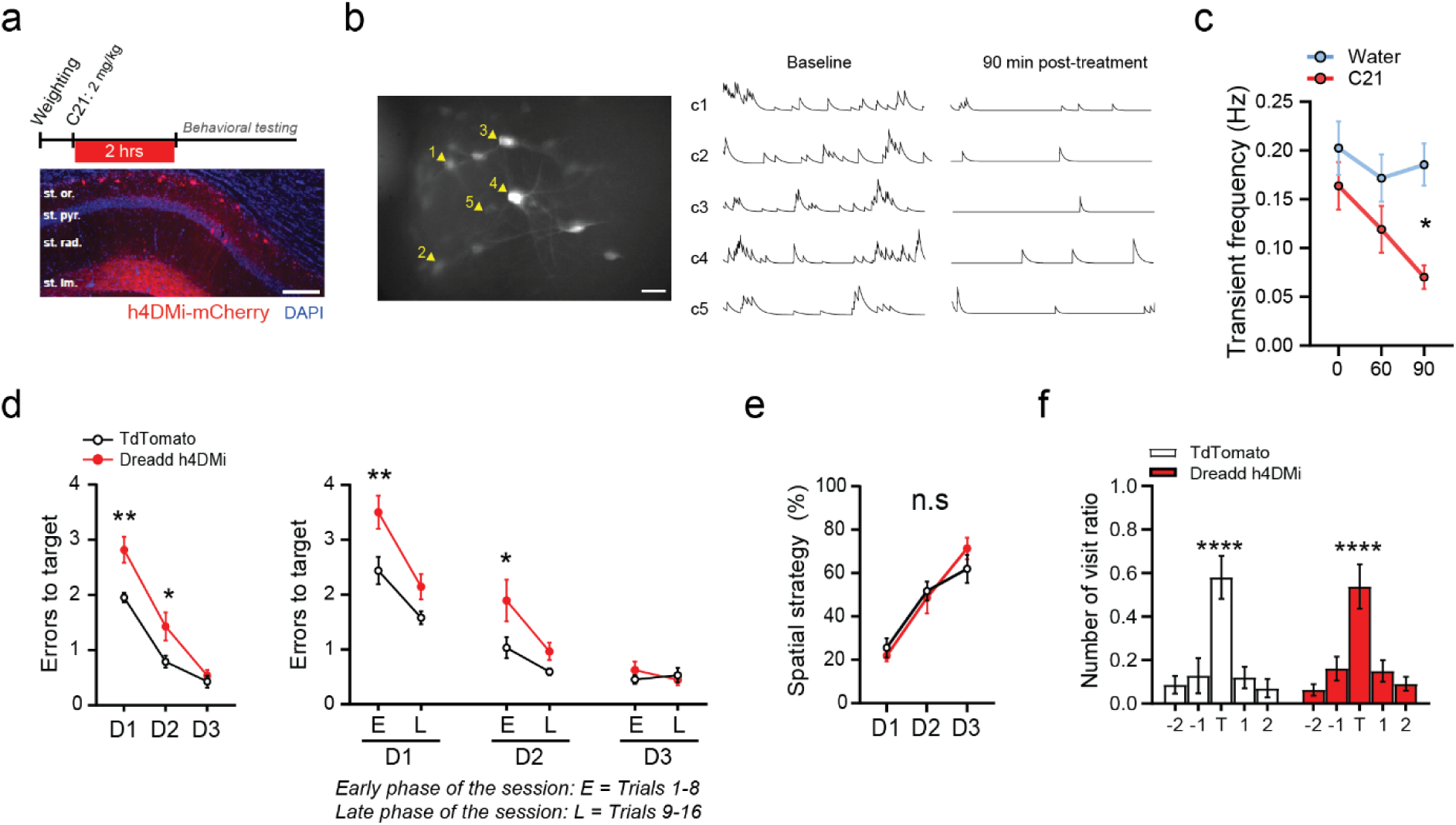
Chemogenetic inhibition of dCA1 O-LM interneurons alter spatial memory formation. **a,** Chemogenetic treatment strategy (top). The DREADD receptor agonist, Compound 21 (C21), was provided 2 hours before behavioral testing. Representative transfection of the DREADD receptor hM4Di (coupled to mCherry) in O-LM interneurons (bottom). Scale bar, 100 µm. **b,** Left: maximum projection of the ΔF/F signal from a representative field of view. Scale bar, 30 µm. Arrows indicate five representative O-LM cells whose calcium traces are shown in panel **(b,** right). Right: ΔF/F traces from the indicated O-LM interneurons during baseline recording and 90 minutes after C21 treatment. **c,** Mean frequency of calcium transients recorded in animals treated with either C21 (n = 10 cells) or water (n = 9 cells). **d,** Left: Day-to-day changes in learning performance: Two-way ANOVA for the effect of C21, session, and their interaction: C21: F(1,13) = 12.58, p = 0.0036; session: F(2,26) = 94.04, p < 0.0001; interaction: F(2,26) = 3.803, p = 0.0356. Right: Day-to-day changes in learning performance, analyzed in blocks of 8 trials (early learning: trials 1–8; late learning: trials 9–16): Two-way ANOVA for the effect of C21, trials blocks and their interaction: C21: F(1,13) = 9.390, p = 0.0090; blocks: F(5,65) = 60.00, p < 0.0001; interaction: F(5,65) = 2.979, p = 0.0175. **e,** Percent of spatial strategies was not affected by O-LM silencing. Two-way ANOVA: C21: F(1,13) = 12.580, p = 0.0036; session: F(2,26) = 94.04, p < 0.0001; interaction: F(2,26) = 3.803, p = 0.0356. **f,** Retrieval memory performance tested on Day 4, showing the number of visits to Star maze arms during 2 minutes of free exploration (no C21 treatment). A t-test against chance level (0.2) yielded **** p < 0.0001. **Statistics:** Panels **d-e,** were analyzed via two-way ANOVA followed by Bonferroni’s multiple comparisons (* p < 0.05, ** p < 0.01). Panel **f** used a one-sample t-test against chance (**** p < 0.0001). Data are presented as mean ± SEM.

## Discussion

Despite evidence that local and distal reference frames influence hippocampal principal networks, including place cells, their impact on hippocampal interneurons has not been determined. Here, by recording calcium activity in freely behaving mice, we reveal how interneuron functions adapt to the type of reference frames being preferentially processed during navigation. Specifically, we found that CA1 O-LM interneurons undergo a dramatic shift in activity depending on the selected frame of reference, whereas VIP interneurons display only minor changes, reflecting the timescale of familiarization processes.

Discrepancies in how neural networks engage during navigation may arise from differences in the type of environmental information they encode, local *versus* distal cues. Comparing these strategies poses methodological challenges, especially when trying to keep sensory inputs and environmental conditions stable. For example, navigating in light *versus* dark conditions changes the frame of reference but also removes key speed cues derived from optic flow^69,70^. To address these issues, we employed *in vivo* calcium imaging with the UCLA miniscope, capturing the natural processing of idiothetic (vestibular self-motion), local (optic flow), and distal (external landmarks) cues, all essential for speed and spatial coding. Prior research shows that identifying the starting location before navigation is critical for relying on a distal frame of reference^52–55^. Accordingly, in the Star maze spatial reference memory task, we developed two paradigms that allow controlling how much mice can discriminate the location of the trial start. When mice had continuous access to distal cues (SA-DC), they swiftly adopted optimized, direct spatial routes, accompanied by higher O-LM calcium activity and more recruited O-LM interneurons by the start arm. By contrast, when mice were deprived of distal cues in the start arm, being enclosed in a box (SA-LC), learning was hindered, leading to inefficient navigation dominated by a local frame of reference. This approach resulted in a serial strategy of sequentially visiting adjacent arms, possibly involving striatal-based associative learning or vestibular-reward associations^71^. Under SA-LC, O-LM cell activity dropped markedly during the start-arm location phase, emphasizing the importance of this brief yet critical period for processing distal information. Notably, both groups received identical environmental information after leaving the start arm, yet SA-LC mice remained unable to match the efficient performance of SA-DC mice. This underscores that distal information gathered during the starting location phase is crucial for effective navigation based on a distal reference frame. Even though this phase accounts for only a small fraction of the total trial duration, it is nonetheless pivotal for successful navigation.

The commonly attributed function of hippocampal interneurons, including SOM cells, is their role in encoding running speed. Speed-related information is considered essential for updating spatial representations during locomotion^27,28,33^ and is often regarded as context-invariant, remaining stable across different environments and over time. SOM interneurons encompass a heterogeneous range of subtypes that are frequently studied collectively as a single group. Among these, O-LM interneurons are especially well-positioned to influence spatial memory, sensory integration, and synaptic plasticity through dendritic modulation of CA3 and entorhinal inputs^36–39^. However, one challenge in studying dorsal CA1 O-LM interneurons via imaging techniques is their sparse distribution and low density, which can result in only a small fraction of such cells being included in broader SOM interneuron datasets. Consequently, any distinct O-LM function may be overshadowed when analyses are averaged across an entire SOM population. By employing the Chrna2-cre mouse line, we specifically recorded O-LM interneurons^36,72^, unlike earlier studies that relied on non-specific calcium imaging^42^, SOM-cre imaging^32,35,43^, or firing-pattern based classification^44^.

Strikingly, when mice navigated using a local frame of reference, a higher proportion of O-LM cells were preferentially tuned during locomotion alongside neutral (locomotion-and immobility-active) interneurons. No immobility-tuned O-LM interneurons were found in those local cues conditions. Furthermore, O-LM cells did not disengage during training; instead, they increased their firing rate and became positively correlated with running speed. Supporting evidence comes from juxtacellular^40,41^ and calcium imaging studies in head-fixed mice^42,32,43,35^ showing enhanced O-LM activity during locomotion in virtual mazes, tactile-enriched running belts, or in freely behaving animals without specific tasks, setups that probably favor local cues to varying degrees. In addition, we further revealed that O-LM interneurons used a rate-coding mode for speed in the Curved-track, the closed arms of the elevated plus maze, and the SA-LC condition of the Star maze, all of which are contexts dominated by local information. In contrast, under a distal frame of reference (e.g., in the SA-DC condition of the Star maze and in the open arms of the EPM), equivalent proportions of O-LM cells exhibited preferential tuning for immobility or locomotion. Notably, their activity became negatively correlated with running speed, shifting from a rate-coding to a population-coding mode. This finding is consistent with the only other study in freely moving rats, on linear tracks and open fields with distal cues access, that reported CA1 O-LM cells showing negative correlation with speed^44^. Another key observation was that, in the SA-LC paradigm, O-LM interneurons reached a peak of activity near the reward location, mirroring the behavior of hippocampal somatostatin-positive interneurons on a local cue-enriched belt^35^. Although VIP interneurons are also regarded as speed-modulated cells^68^, we found that their speed-encoding properties remain unaffected by the frame of reference. Across all conditions, VIP interneurons primarily displayed population-coding for speed, showing only a slight firing-rate increase during locomotion on the first day of exposure in the Curved-track and SA-LC conditions, an effect consistent with novelty detection^59^, which can extend into longer familiarization (Curved track, Day 2–3).

Crucially, by using O-LM (but not VIP) speed scores, we could decode the animal’s navigation strategy, suggesting that O-LM interneurons are promising candidates for understanding how navigation strategies drive hippocampal engagement in spatial information processing. Our study thus provides new insights into interneuron speed encoding by showing that: (i) while many hippocampal interneurons are speed-tuned, not all adjust their encoding depending on navigation strategy; (ii) for O-LM interneurons, the sign of the speed correlation (positive or negative) reflects how the animal processes environmental cues, with negative correlations linked to a dominant distal frame of reference and successful spatial memory; and (iii) experimental conditions balancing different frames of reference, such as the elevated plus maze or our two Star maze paradigms, are necessary to fully understand hippocampal cell’s functions.

It is well established that place cells rely on both local and distal cues^2^. Although CA1 interneurons can also exhibit spatial tuning^35,60–63^, the effect of different reference frames on their place fields remains unclear. Here, we found that the place fields of O-LM and VIP interneurons are influenced by whether mice navigate using a distal or local frame of reference. First, O-LM interneurons exhibited more place fields on average when navigation was driven by local cues (e.g., the curved track and SA-LC, Day 1–2), whereas VIP place cells remained relatively stable regardless of context or strategy. Second, O-LM place fields were primarily located at decision points under distal-based navigation but shifted to the inner alleys under local-based navigation. Similarly, VIP place fields moved from the rewarded target arm (in distal-based navigation) to non-target arms (in local-based navigation), paralleling findings by Turi and colleagues^65^ that CA1 stratum pyramidale VIP interneurons are modulated by reward expectation. This shift likely reflects the difficulty in pinpointing reward locations under a local reference frame, as mice may regard each arm as a potential reward source. Third, the number of place fields changed with re-exposure to the environment in a context-dependent manner. Although the proportion of VIP place fields remained relatively stable, O-LM place fields in the Star maze dropped from around 20% to 10% when mice habituated to local-based navigation. This decrease aligns with the reported proportion of place cells among SOM-positive CA1 interneurons after extensive training^35^ and cannot be explained by a general disengagement of O-LM cells, as their firing rate increased with training. Hence, both the dominant reference frame and repeated exposure to the task environment can significantly shape interneuron place field properties.

The Star maze is a complex, multi-choice environment involving distinct learning phases, location, navigation, reward consumption, post-reward, and transportation, each likely engaging neural circuits differently during spatial reference memory formation. Focusing on O-LM interneurons, we found that their activity in the start arm location phase depends on the type of information available (local *versus* distal), their place fields and speed encoding vary with the selected navigational frame of reference, and their reactivity near the reward zone peaks when navigation relies on local cues. To assess the overall impact of O-LM inhibition in dorsal CA1, we used chemogenetics (DREADDs) under the SA-DC paradigm of the Star maze. Our results align with previous findings showing that O-LM silencing can affect memory function^38,73^. Although these cells constitute a subset of somatostatin (SOM) interneurons and are sparsely distributed in CA1, we found that they are crucial for optimizing spatial reference memory formation, particularly in the early phases of training (first eight trials on Days 1 and 2), a period that coincides with the peak O-LM firing rates. However, inhibiting O-LM cells did not impede the use of a distal reference frame or completely disrupted spatial learning and memory as observed on Day 3, suggesting that alternative circuits, potentially involving other brain regions may compensate and favor the use of a distal reference frame. It should be noted that hippocampal mechanical lesions need to extend to around 20% of the dorsal hippocampus to induce dramatic spatial learning impairments in Morris Water-Maze^83^. Here, we targeted specifically O-LM interneurons, and this only in the dorsal CA1 subregion, so silencing the entire O-LM network across the hippocampal axis could elicit more pronounced memory impairments.

CA1 O-LM interneuron activation critically modulates principal network activity by facilitating CA3 Schaffer collateral inputs while inhibiting entorhinal temporoammonic (TA) inputs^36,39^. CA3 itself is crucial for long-term memory storage and retrieval, as well as for creating unified temporal and spatial contextual representations that support CA1 computations involved in learning^74–79^. CA3 is especially active during immobility^80^, and its Schaffer collateral inputs are strongest when animals are immobile or moving slowly^76,81^. We observed enhanced O-LM engagement during the start-arm location phase under distal cues, when mice were largely immobile or moving slowly. Identifying each unique start point may require distinct spatial engrams in CA1 neurons and their stabilization over time, a process potentially supported by robust CA3-to-CA1 information transfer. Consistent with this idea, increased O-LM activity promotes place cell stability, whereas reduced O-LM activity facilitates plasticity and remapping^37,82^. Accordingly, in the SA-DC condition (distal frame of reference), heightened O-LM activation during start-arm identification may help form stable place cell engrams. During the Star maze navigation phase, O-LM firing rates were significantly higher during locomotion in the SA-LC condition (local frame of reference) compared to the SA-DC condition. Building on Udakis and colleagues’ proposal, this elevated O-LM activity could convey the hippocampal system’s effort to establish a stable representation of the local environment (e.g., maze geometry). Ultimately, this may have contributed to the modest improvement in navigation performance observed across the three learning days.

Taken together, this work demonstrates that O-LM interneurons exhibit substantial changes in firing patterns, speed encoding, and place-field organization depending on whether mice navigate using local or distal reference frames, whereas VIP interneurons remain relatively stable. These findings underscore a distinct role for O-LM cells in orchestrating context-dependent hippocampal function, revealing their pivotal contribution to the early optimization of spatial reference memory. This illustrates that hippocampal interneurons can be more flexible in their functions than generally presumed, mirroring the flexible coding abilities of principal cells^88^. This work also suggests that the reference frame should be more systematically investigated to fully grasp the function of hippocampal networks as shifts in the reference frame could have dramatical effects on hippocampal neuron functions. This would ultimately aid the refinement of models of hippocampal functions.

## Methods

### Animals

All procedures were carried out following the recommendations of the Canadian Council on Animal Care (protocol 2015-7650) and in agreement with approved procedures by McGill University Animal Care Committee. A total of 44 male mice, aged 6 to 7 months, were used in this study. Chrna2-cre mice, which are on a C57BL6 genetic background, were specifically utilized to target O-LM interneurons^36^. These mice were crossbred with tdTomato reporter mice (tdTomato: B6;129S6-Gt(ROSA)26Sor tm9(CAG-tdTomato)Hze/J, #007905; Madisen et al., 2010) to generate Chrna2-tdTomato mice, in which tdTomato is exclusively expressed in cells containing Cre recombinase. The tdTomato signal was used to confirm the presence of intact O-LM interneurons during baseplate fixation, as the GCaMP6f calcium signal was significantly reduced under anesthesia. Two groups of mice were formed for the study: one with 17 Chrna2-cre mice for calcium imaging and another with 19 mice for chemogenetic experiments. Heterozygous VIP-cre mice were produced by crossing homozygous Vip-IRES-cre (VIPcre, #010908, The Jackson Laboratory) with homozygous Ai9 lox-stop-lox-tdTomato cre-reporter mice (#007905, The Jackson Laboratory). For the study of VIP interneurons, 8 mice were used. Both mouse lines were housed in groups of four to five per cage and were subsequently individualized following the first surgical procedure at the age of 5 to 6 months. The animals were maintained on a 12-hour light/dark cycle at 22°C and 40% humidity, with food and water available *ad libitum*. All experiments were conducted during the light phase.

### Surgical procedures

For all surgical procedures, mice were first anesthetized in a chamber filled with a mix of oxygen and 5% Isoflurane before being transferred to a stereotaxic frame (David Kopf Instruments), where mice were supplied with 2.5-2% isoflurane in oxygen until the end of the procedure. For reducing pain and inflammation as well as for fluid replacement, Carprofen into saline was subcutaneously administered before and at the end of the surgery as well as postoperatively for 3 days. Eyes were lubricated with an ophthalmic ointment (Optixcare) and body temperature was maintained using a heating pad. The skull was cleared of connective tissues. A dental drill was used to perform craniotomies for injection and lens implantation.

### Viral injections

For miniscope calcium imaging experiments, the adeno-associated virus (AAV) expressing the GCaMP6f calcium reporter AAV2/9.Syn.GCaMP6f.WPRE.SV40 (Penn Vector Core, ≥ 7×10¹² vg/mL) and AAV9.CAG.Flex.GCamp6f.WPRE.SV40 (Penn Vector Core, 5.248e13 GC/ml) were used to image OLM and VIP calcium activity, respectively. Both viruses were diluted in sterile saline (1:4). For DREADD chemogenetic inhibition of OLM interneurons, the cre-dependent AAV2-hSyn-DIO-hM4D(Gi)-mCherry (Addgene #44362, 5×10^12^ vg/ml) or control Tdtomato expressing virus AAV2-Ef1α-flex-tdTomato (Laval-Canadian Neurophotonics platf., 4.0e12 GC/ml, no dilution) were injected into the dorsal CA1 from Chrna2-cre mice. Briefly, after exposure of the skull, a small craniotomy was performed above the dorsal CA1 region (AP: -1.9, ML: ±1.4, DV: -1.3). The virus suspension was infused at a rate of 1 nl/sec using pulled glass pipettes connected to a nanoinjector (Nanoject III, Drummond Scientific, Broomall, PA). For calcium imaging experiments, 700nl of the virus was unilaterally injected into the right hemisphere. For chemogenetic experiments, 500 nl (undiluted) of DREADD-hM4Di or control virus was bilaterally injected. Pipettes were left in place for 10 min before being carefully withdrawn. The skin was then sutured and protected using Polysporin to control for potential post-surgical infection. Following the surgery, animals were closely monitored for 3 Days, with daily s.c. injections of Carprofen.

### GRIN lens implantation

Two to three weeks post-injection, mice were implanted with a 1.8 mm diameter Gradient Refractive Index lens (GRIN; Edmund Optics), just above the dorsal CA1 region. Briefly, craniotomy was performed by drilling a 2 mm diameter hole in the skull: -2.2 mm anteroposterior and centered to 1.8 mm lateral from the bregma. Bones fragments and dura were carefully removed with small sterile tweezers. 27 Gauge blunt tip needles were used to gently aspirate the cortical tissue until striations of the corpus callosum were visible and 30 Gauge needles were used to carefully expose the alveus. Brain tissue was moistened with sterile saline all along the digging process. The GRIN lens was implanted on top of the alveus fibers (depth ∼-1.30 mm) and secured to the skull using dental cement (C&B-Metabond, Patterson Dentals). A small cap embedded into surgical silicone adhesive (Kwik-Sil ^TM^; World Precision Instruments) was employed to protect the GRIN lens until the next surgery. 5-6 weeks after lens implantation, an aluminum baseplate was mounted on the animal’s head to secure the miniscope for calcium recordings. The baseplate was fixed to the skull at the optimal focal distance via dental cement and protected with a custom 3D-printed plastic cap during non-recording periods in their home cage.

### Calcium & behavior recordings

*In vivo* calcium imaging was conducted using a one photon UCLA miniscope (v3; miniscope.org), while simultaneously recording behavioral data. This was accomplished with the Miniscope acquisition software (miniscope.org), utilizing a consumer-grade webcam (Logitech) positioned 50-100 cm above the behavioral apparatus. The data acquisition (DAQ) box was connected via a USB host controller (Cypress, CYUSB3013) and linked to a flexible coaxial cable for data collection. The miniscopes were assembled according to open-source instructions found at miniscope.org. The DAQ collected behavioral data at 30 Hz and cellular imaging data at 15 and 20 Hz to capture calcium transients from O-LM and VIP interneurons, respectively. Recordings were saved as uncompressed .avi files containing 1000 frames, paired with corresponding timestamps to facilitate precise temporal alignment of the behavioral and calcium imaging data during post-hoc analysis. To minimize the risk of photobleaching while ensuring good signal quality for further analysis, the miniscope excitation LED power was set to between 5% and 10%.

### Behavioral procedures

#### Habituation prior behavioral testing

Before behavioral testing, the mice were gently handled for 2 to 4 minutes each day over a period of 5 to 6 days to help them become accustomed to the experimenter. Following this, they were habituated to the plugging and unplugging of the miniscope, as well as the experimental room, for an additional 2 days.

#### DREADD inhibition

To minimize potential stress from repetitive intraperitoneal (i.p.) injections, we created an experimental condition that encourages spontaneous absorption of the DREADD ligand Compound 21 (C21, 2 mg/kg) mixed in 10% sucrose water. Briefly, an empty housing cage (neutral environment) was equipped with a small water port (100 µl capacity). The animals had free access to the water port and were removed from the environment once they consumed the drug. To reduce neophobia, the animals were given 10% sucrose water for two days before starting the C21 treatment. To examine the pharmacokinetic properties of C21 following oral absorption at a dose of 2 mg/kg, Chrna2-cre mice were co-injected with a mix of DREADD inhibitory AAV2-hSyn-DIO-hM4D(Gi)-mCherry virus or a control TdTomato-expressing virus. Both viruses were combined with AAV2/9-Syn-Flex-GCaMP6f. The mice had the opportunity to visit a neutral environment every 30 minutes. O-LM calcium activity was recorded for 5 minutes during each exposure session, with the mice returning to their home cage between sessions. Following analysis of calcium transients frequencies (**Fig. 8a-c**), final DREADD experiments were performed with experimental mice that were provided with C21 two hours before testing. During the two-hour waiting period prior to testing, the animals remained in their home cage.

#### Behavioral testing procedure

For each behavioral task, the behavioral devices were thoroughly cleaned between mice and sessions using Peroxigard. Except for the Elevated Plus Maze experiments, the mice were subjected to water restriction for two days before any behavioral procedures that required it. During the water restriction periods, mice were provided with 40% of their average daily water intake during testing. After the tests, they were given two hours of free access to water in their home cages.

#### Elevated Plus Maze

The apparatus was constructed of plastic and consisted of two open arms and two closed arms (40 cm long), positioned opposite each other and connected by a central area. The closed arms were surrounded by 20 cm high opaque black walls. The maze was elevated 60 cm above ground. The experimental room was illuminated with a ∼ 650 light lux. The testing began by placing the animal in the center of the maze, facing one of the enclosed arms. Mice were then allowed to freely explore the maze for 5 minutes. The percentage of time spent in open *versus* closed arms^84^, the mean speed, and total distance were measured to evaluate the anxiolytic-like response. Additionally, the percentage of locomotion and immobility time were also measured.

#### Curved track

Water-restricted mice were trained to run on a custom curved track designed to prevent them from having direct visual access to the reward zones. To eliminate access to distant cues, 16.3 cm high opaque white walls enclosed the maze. This design concealed the experimenter’s hand movements and encouraged the mice to focus more on their immediate surroundings while navigating the track. The apparatus was 140 cm long and made of white plastic. Reward zones, where the mice received water, were located at each end of the track, positioned 5 cm from the extremities. Each session started with the introduction of the mouse into the right end of the track blocked by a door. After approximately 10 seconds, the door was open, allowing the mouse to run the track to obtain 25 µl of 10% sucrose water for each visit to one of the two reward ports. The mice completed 28 laps per session over three consecutive days. The percentage of time spent in locomotion, the average speed, total distance traveled, and percentages of locomotion and immobility were recorded.

#### Star Maze task

The star Maze apparatus consisted of five arms (5 cm large and 20 cm long) radiating from a central ring made of five pentagonal segments (5 cm large and 20 cm long). These segments were connected via triangular choice points (**Fig. 1**). The maze was constructed from white plastic. Each arm’s extremity was equipped with a tube (diameter 1 cm) connected to a foot pump to deliver air puff. Air puffs were provided in non-target arms only to motivate the animal to escape and encourage them to learn to avoid non-rewarded paths. We developed a spatial reference memory version of the original star Maze task^56^. Briefly, water restricted mice had to learn the location of a target arm, where 50 µl of 10% sucrose reward was delivered during each trial. The mice were trained during 16 trials per day over three consecutive days (learning phase). Memory retrieval performances were tested on Day 4 (probe trial) that lasted 2 minutes and did not involve water reinforcement. Each trial started by placing the animal in one of the four possible start arms, which was closed off by a door. The order of the star-arm locations was pseudo-randomly determined to prevent the same sequence from repeating and to avoid the development of egocentric strategies. After approximately 20 secs spent in the start arm (location phase), the door opened, allowing the mice to navigate the maze (navigation phase) until they found the reward. Two distinct procedures were used to manipulate access to distal cues during the start-arm phase. In the SA-DC procedure, animals were transported from the target arm to a new start arm location using a square platform (5 × 5 cm) with 1 cm high edges. In this protocol, mice had access to distal cues during both transport and within the start arm. Conversely, in the SA-LC procedure, animals were transported in a closed box and remained there throughout the start arm location period, which eliminated access to distal cues in those phases. During the learning phase (Day 1 to Day 3), the number of errors (entries into a non-target arm), the percentage of trial utilizing spatial strategy (direct trajectory), the mean speed and the percentage of immobility *versus.* locomotion was measured to assess the efficiency of spatial memory formation. During the probe test on Day 4, the arm visit ratio was measured to evaluate retrieval performance.

### Neural Data Analysis

MATLAB (MathWorks) and Graphpad Prism were used for all data analysis.

#### Video preprocessing & calcium and behavior data synchronization

The raw video resolution was 752 by 480 pixels. Videos were down sampled in space (3x using bi-linear interpolation). Preprocessing was made with MATLAB functions, NorMCorre for motion artifact correction^85^ and CNMFe for cell segmentation and calcium transients extraction^86,87^. Behavioral videos were acquired at 30 frames/seconds, and matched with calcium data using miniscope software timestamps files.

#### Behavioral tracking

To extract information about the mice’s position and running speed, we used a custom video-tracking function written on Matlab 2018b. Briefly, the mouse’s body was detected using background image subtraction, and the body’s center of mass was used for the tracking process. The spatial location data were interpolated to match the calcium imaging sampling frequency using linear interpolation. Running speed was calculated by computing *Δd/Δt* where *d* is the distance between two subsequent frames and *t* the time between those frames. To eliminate detection artifacts, we applied a Gaussian filter with a sigma of 66 ms to smooth the results. The resulting XY position data and speed signals were then plotted for each mouse and each session for visual inspection to confirm good signal quality.

#### Calcium signal extraction

Calcium imaging data analysis was conducted using MATLAB 2018b with custom script and functions. Initially, rigid motion correction was applied using NorMCorre (https://github.com/flatironinstitute/NoRMCorre). Calcium ΔF/F traces were then extracted using CNMFe algorithm with the following parameters: gSig = 3 pixels (width of gaussian kernel), gSiz = 20 pixels (neuron diameter), background model = ‘ring’, spatial algorithm = ‘hals’, min corr = 0.8 (minimum pixel correlation threshold), min PNR = 8 (minimal peak-to-noise ratio threshold). Raw calcium traces were filtered to remove both high-frequency fluctuations and slow drift prior to binarization. Neurons were considered active when normalized calcium signal amplitude exceeded two standard deviations and the first-order derivative was above 0. The final binarized rising phase vector was then set to 1 whenever when neurons were active, and 0 otherwise. This binary vector was treated as the firing rate in all further analyses.

#### Locomotion and immobility analysis

The running speed signal was used to identify periods of locomotion (defined as >5 cm/sec) and immobility (defined as <5cm/sec). This distinction allowed for the analysis of modulation in O-LM and VIP calcium activity across these states. Additionally, both locomotion and immobility periods had to last for at least 1 sec to be included in analysis. To extract cell tuning, the average frequency of activity during each state was computed and then compared to a random distribution generated by 1000 iterations of time-shuffling. For each iteration, a random frame was selected to split the trace into two bouts, which were then permuted to un-correlate the calcium signal from behavioral data while preserving the temporal structure of calcium activity. For a cell to be classified as significantly preferring either locomotion or immobility, its activity must be greater for the preferred state and exceed the 99th percentile of the shuffled distribution.

#### Place field analysis

Place fields were computed from binarized calcium signals^64^. Analyses were conducted on the binary vector of the rising phases of calcium transients as if it were the firing rate of the cell. Rate maps were constructed by binning the position data into pixels corresponding to a 2cm×2cm grid of locations. To be classified as a place cell, neuron’s within-session split-half rate map correlation had to exceed the 95th percentile of a shuffled control distribution. The shuffled control was computed for each cell by randomly circularly shifting its firing rate vector relative to the position data by at least 30s and recomputing the split-half rate map correlation. This process was repeated 1000 times to create the null distribution. This classification allowed only to consider as putative place cells neurons that show stability in their spatial rate-maps. We then computed for each of those candidate place cells the existence of at least one place field: spatial bins in which the activity was higher than the 95th percentile of a shuffled control distribution were considered as part of a place field. Place fields were defined by grouping spatial bins with significant activity that were adjacent. Place fields locations were then extracted by computing the centroid location of the place fields.

#### Speed analysis

Cell speed tuning was assessed by calculating a speed score for each interneuron. Speed was binned into 2cm/s bins and the average activity of each cell was computed for each speed bin. Speed score was defined as the Pearson’s correlation coefficient between a cell activity and the mouse speed. Due to the limited number of O-LM interneurons per mouse, all cells from all mice were grouped in a pseudo-population to generate population vectors (PV) of the running speed that provide a representation of how the O-LM population as a whole encodes each speed bins. To generate the speed population vector matrix, the speed firing rate distributions of all recorded interneurons from a given condition (or maze region type) were combined into a single, two-dimensional array, with cell number in the rows and speed bins in the columns. Each column of this matrix represents an estimate of the composite population vector for the corresponding running speed.

#### Preferred activation phase in Star-maze

To determine whether O-LM and VIP interneurons had a preferred phase of activation in the Star-maze task, the average frequency of calcium transients of each cell detected was computed for the main behavioral phase. These phases included the start arms periods (by location relative to the target), the active navigation (by locomotion state: immobility, locomotion to errors & locomotion to the target), the reward consumption and post-reward exploration (by locomotion states). For each interneuron, we compared the recorded activity to a null distribution created from 1,000 iterations of time-shuffling. Activity levels exceeding the 99th percentile of the surrogate distribution was considered significant. Subsequently, all recorded cells from each condition of the star maze were grouped into a pseudo-population to compute the population vector (PV), which illustrates how the overall O-LM and VIP interneuron populations encode each behavioral phase of the task.

#### Sampling size matching for Elevated Plus Maze

As the mice spent more time in the closed than in the open arms of the EPM, we resampled the data to ensure that differences between the two arms were not due to a smaller sampling size for open arms data. For each mouse we extracted the total time spent in open arms and used that value as a duration threshold. This allowed us to restrict the amount of closed arms data included to match the sample sizes between both arm types.

#### Decimation analysis for Star-maze

As the total number of O-LM and VIP-pyr interneurons recorded in the “SA No Distal Cue” condition was lower than in the “SA Distal Cue” condition, we run a decimation analysis on “SA Distal Cue” data to evaluate if the results from “SA No Distal Cue” condition could be due to the reduced sampling size. For each “SA Distal Cue” star-maze session, we randomly selected the same number of O-LM as in the “ SA No Distal Cue” condition and recomputed the results. This process was repeated for 1000 iterations. The resulting distribution was used to compute a 99% confidence interval for start-arm PETH from “SA Distal Cue” condition (**Fig. 2h**). Additionally, for each decimation iteration, we ran a correlation with each original dataset to determine if decimated traces from “SA Distal Cue” were more similar to the original “SA Distal Cue” traces or to the “SA No Distal Cue” traces.

#### SVM decoding of the reference frame

To decode the reference frame from the interneurons speed scores, we used a SVM algorithm implemented in MATLAB (*fitcsvm* function) trained for two-class classification of the reference frame. The predictor data to which the SVM classifier was trained and provided to the model for decoding was the average speed-score per session, as the number and identity of cells vary across sessions. Predictor data were standardized using the weighted standardizing algorithms inbuilt within the *fitcsvm* MATLAB function. The class labels were either proximal (closed arm of the EPM, linear track, SA-proximal cues condition of the Star-maze) or distal (open arm of the EPM, SA-distal cues condition of the Star-maze). To avoid overfitting, data to decode were removed from the training dataset used to generate the classification model.

### Statistical analysis

All data are presented as mean ± standard error to the mean (SEM) and statistical test details are described in the figure legends. The normality of each group distribution was assessed using the Shapiro-Wilk normality test before using the parametric test. Graphpad prism software was used to run One-way, two-way or repeated-measure ANOVA to determine significance of the effects before running post-hoc Sidak test (accounting for multiple comparison) to localize the effect.

### Histology and immunochemical staining

#### Validation of GRIN lens placement

After completing the experimental procedures, animals were perfused to verify the GRIN lens position. Mice were deeply anesthetized with intraperitoneal injection of a cocktail of ketamine/xylazine/acepromazide (100, 16, 3 mg/kg, respectively). Following anesthesia, the mice were transcardially perfused with 4% paraformaldehyde in 0.01M phosphate-buffered saline (PBS). The perfused brains were soaked in 4% paraformaldehyde at 4 °C overnight and then transferred to 1× PBS. The dorso-intermediate hippocampus was sliced into coronal sections (50 µm) using a vibratome (plate number 1, Leica Biosystems). Half of the sections were directly mounted on glass slides and coverslipped with Fluoromount-G containing DAPI (Southern Biotechology).

#### Slide scanner imaging

Fluorescent images from the dorsal CA1 region were digitally captured at 10× magnification using an Olympus Slide Scanner (Olympus VS120) with appropriate light excitation wavelengths. The images were then processed using ImageJ software (NIH) to ensure uniform adjustments of brightness and contrast across all images.

### Data and code availability

All data reported in this paper will be publicly available upon manuscript publication in an online database or via request to the corresponding authors. All source codes will be deposited at Github. Further additional information and requests should be addressed to the lead contact, Sylvain Williams.

## Acknowledgments

We thank Dr D. Aharoni and the miniscope.org community for providing help with the V3 miniscope technology. We thank Ke Cui for providing technical assistance with this project. We thank Prof. Mark Brandon for sharing the VIP-cre mouse line and Prof. Klas Kullander for sharing the Chrna2-cre mouse line. We deeply appreciated the valuable inputs provided by Drs Romain Goutagny, Jesse Jackson, Quinn Lee, Guillaume Etter and Pola Tuduri on prior versions of this manuscript.

## Author contributions

Conceptualization was carried out by L.P and J.B.B. Design and validation of the behavioral paradigms Curved track and Star maze was performed by L.P. Investigations were carried out by L.P, J.B.B, S.A, and M.C. The analysis strategies were defined by L.P, and J.B.B. Matlab codes were written by J.B.B and E.G.L. Data analysis was done by J.B.B, L.P and S.A. Funding was acquired by S.W. Supervision was the responsibility of L.P., J.B.B. and S.W. The original draft was written by L.P and J.B.B. Review and editing of the draft was carried out by L.P, J.B.B, and S.W.

## Supplementary Figures

This document includes:

Extended Data Figures 1-12

**Extended Data Fig. 1.**
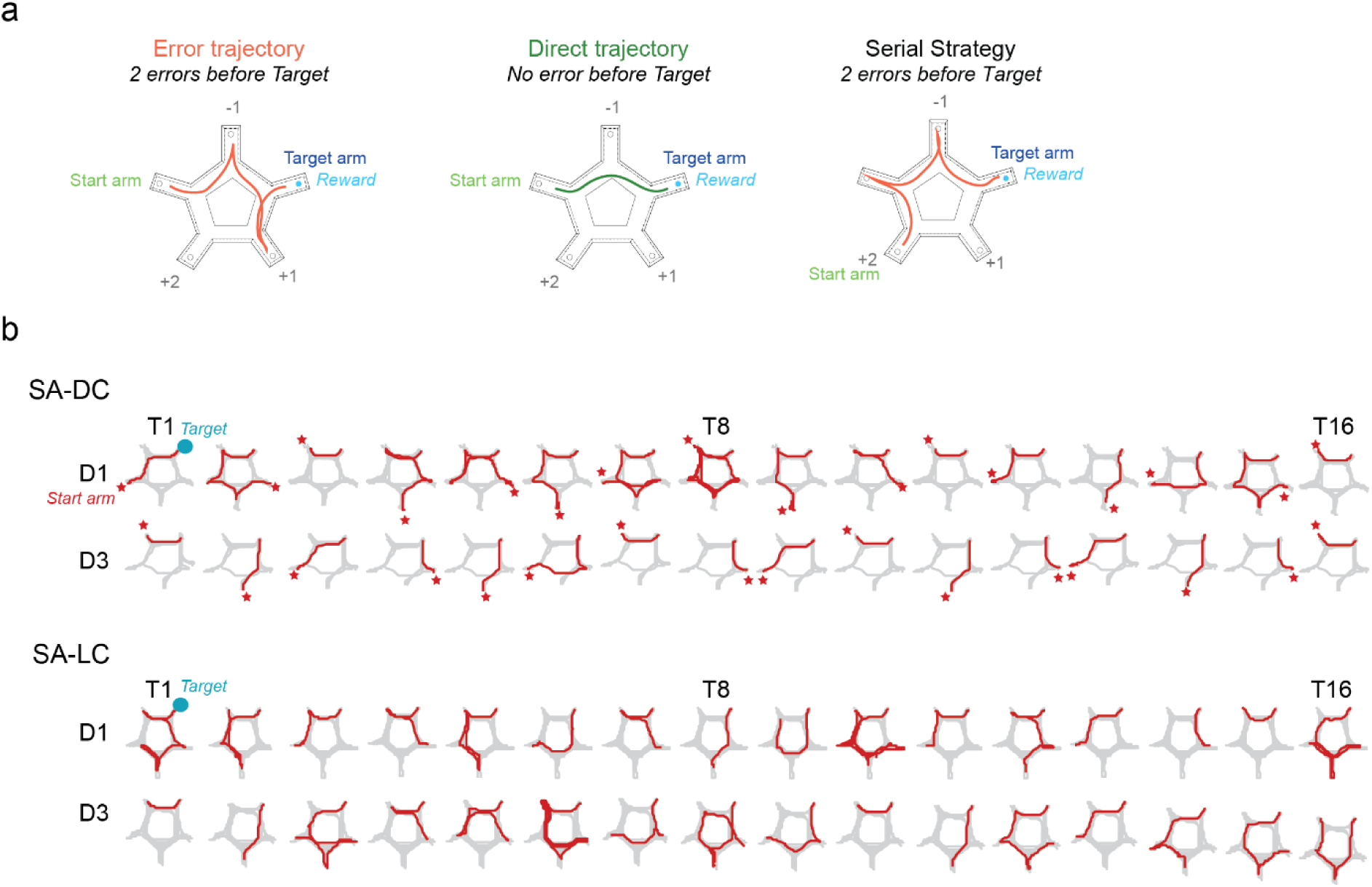
Example of trajectories performed during successful (SA-DC) versus poor (SA-DC) spatial reference memory. **(a)** Schematic representations of a non-direct error trajectory (Left, in red, 2 errors before target) and a trajectory performed directly (Right, in green) from the start (green) toward the target (dark blue) arms. **(b)** Example of representative trajectories for the first (Day 1) and last (Day 3) training sessions (16 trials), for mice tested either in the Star maze conditions SA-DC (Top) or SA-LC (Bottom). The trial trajectories are displayed in red, and all session trajectories are displayed in gray. For each trial, the start arm location is indicated with a red star. These examples are coming from Chrna2-cre mice.

**Extended Data Fig. 2.**
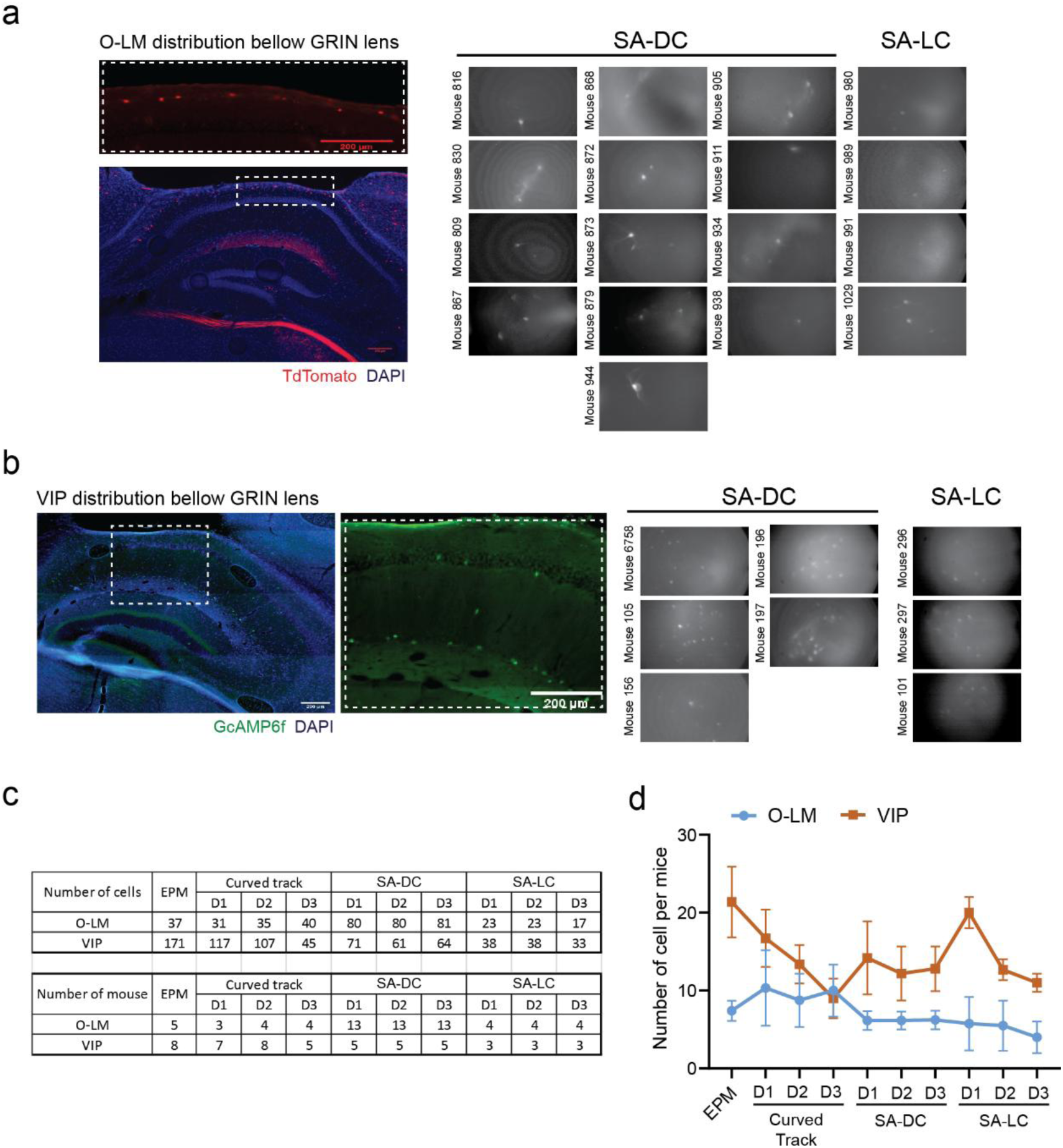
Histological representation of GRIN lens position and max projections representing the field of view selected for each animal. **(a)** Left: magnified view (top) from a representative micrograph showing the distribution of Tdtomato positive O-LM interneurons (bottom). The cells are sparsely distributed below the GRIN lens. As the GcAMP6f signal was low following anesthesia (isoflurane) and perfusion of the animal, the TdTomato signal was used for illustration. Right: maximum projection of the DF/F signal showing the field of view recorded for each Chrna2-cre mouse used in the SA-DC (n = 13) and SA-LC (n = 4) conditions of the Star maze. **(b)** Left: representative micrograph showing the distribution of GcAMP6f-positive VIP interneurons and associated magnified view. In contrast to O-LM interneurons, the VIP cells maintained a good level of GcAMP6f expression after perfusion. Right: maximum projection of the DF/F signal showing the recorded field of view for each mouse used in the two versions of the Star maze (SA-DC, n = 5 and SA-LC, n=3 mice). The blue channel is DAPI. Scale bars: 200 µm. **(c)** Top: table of the total number of cells per experiment. Bottom: table of the number of mice per experiment. **(d)** The average number of cell detected per mice and per session was greater in VIP-cre than in Chrna2-cre mice due to the lower density of O-LM (Mixed-effect ANOVA: session effect F(3.314,29.46)=1.104, p=0.0367; Cell population effect F(1,19)=15.97, p=0.0008; Cell population * session interaction F(2,42)=1.105, p=0.3675; α=0.05).

**Extended Data Fig. 3.**
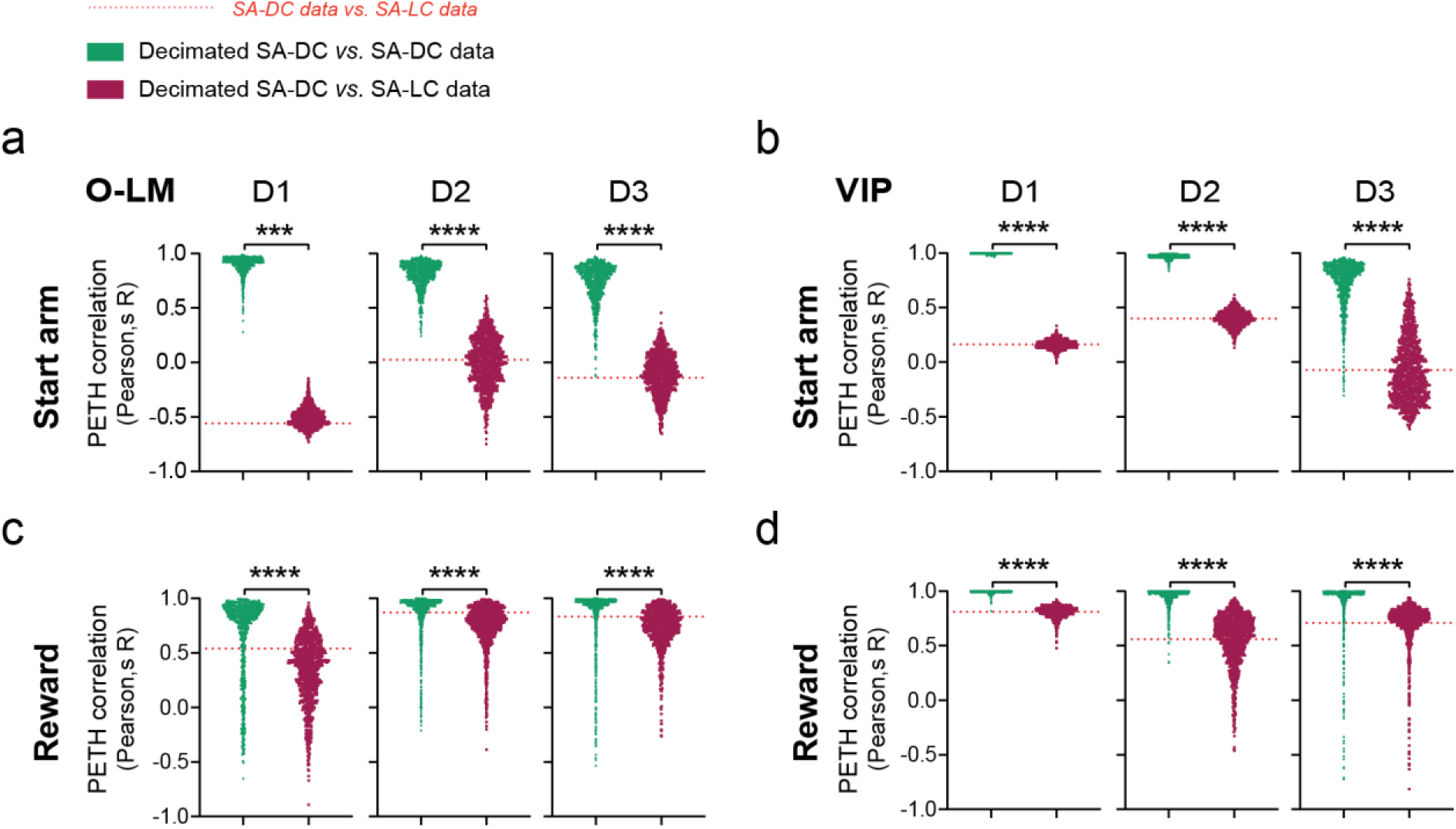
Decimation analysis on the SA-DC dataset confirmed that cell activation during start arm and reward in the SA-LC conditions are unlikely to be due to smaller sampling size. To control for the smaller sampling size effect in SA-LC condition, 1000 iterations of decimation were run: for each iteration i) a number of cells equal to the number of cells from SA-LC condition was randomly selected from the SA-DC dataset and ii) the response profile for the Start arm PETH **(a-b)** and the reward PETH **(c-d)** were computed for those downsampled SA-DC dataset and iii) Pearson’s correlation r value were computed between these downsampled traces and the original SA-DC data (green) and the SA-LC data (purple) to measure the similarity between traces. In addition, the Pearson correlation r value was computed to display the observed similarity level between the original SA-DC trace and the SA-LC trace (red dotted line) as a reference for the decimated results. Each dot represents an individual decimation iteration result. For PETH of averaged DF/F activity round the location phase in the start arm, decimated SA-DC traces were more similar to original SA-DC traces than SA-LC traces for both **(a)** O-LM (Day 1: t(999)=316.3, p<0.0001; Day 2: t(999)=245.8, p<0.0001; Day 3: t(999)=122.5, p<0.0001) and **(b)** VIP (Day 1: t(999)=650.4, p<0.0001; Day 2: t(999)=109.6, p<0.0001; Day 3: t(999)=56.49, p<0.0001). For PETH of averaged DF/F activity round the reward consumption, decimated SA-DC traces were more similar to original SA-DC traces than SA-LC traces for both **(c)** O-LM (Day 1: t(999)=32.71, p<0.0001; Day 2: t(999)=27.42, p<0.0001; Day 3: t(999)=16.54, p<0.0001) and (D) VIP (Day 1: t(999)=129.3, p<0.0001; Day 2: t(999)=61.67, p<0.0001; Day 3: t(999)=54.19, p<0.0001). **Statistics: (a-d)** Differences between conditions on the same day: *** p<0.001 and, **** p<0.0001. Data are shown as mean ± SEM

**Extended Data Fig. 4.**
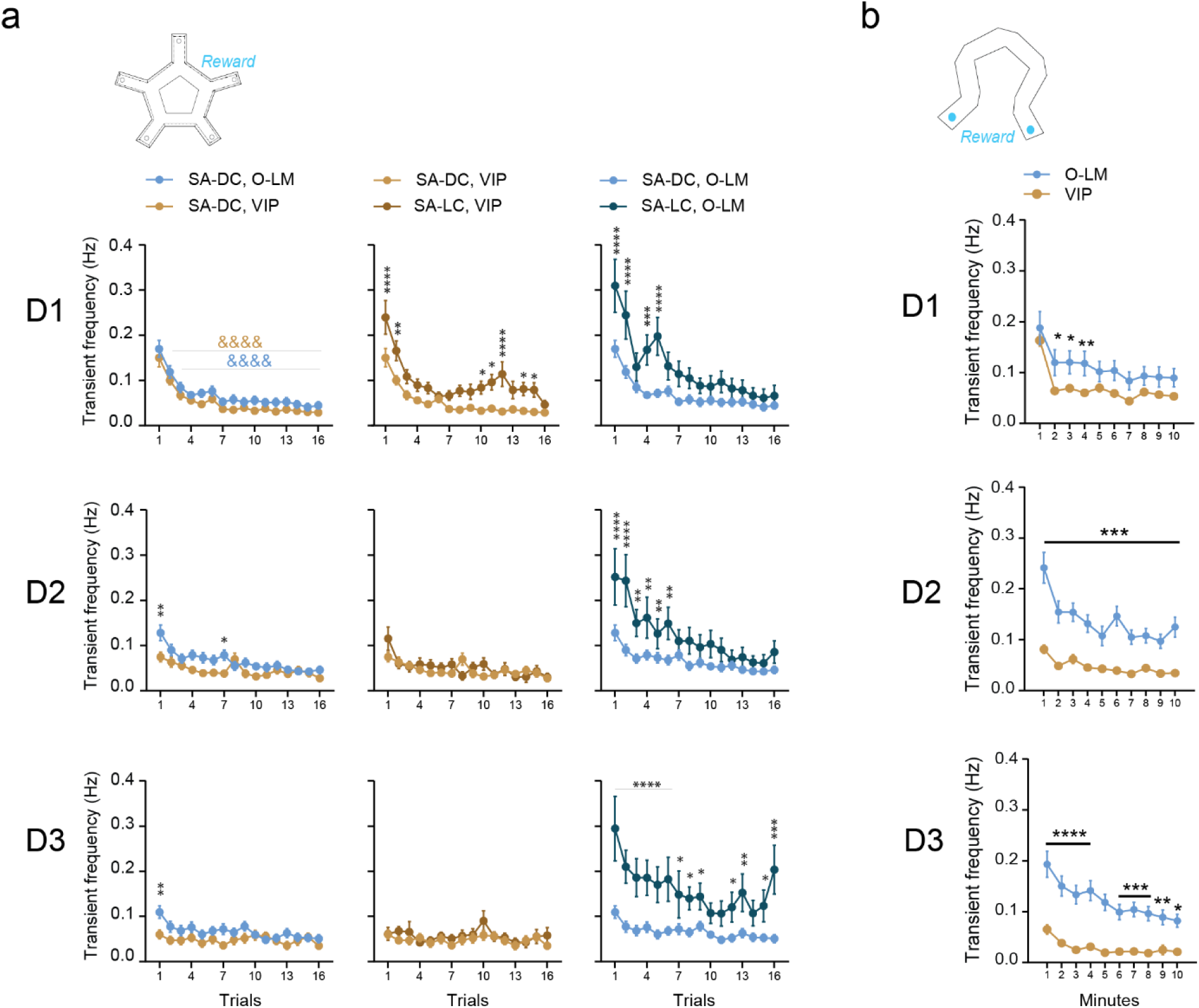
Evolution of O-LM and VIP firing rates during navigation in the Star maze and early exposure of the curved track. **(a)** Activity levels across trials in the Start maze procedure. In the SA-LC condition, activity levels are higher during navigation in both O-LM (Days 1 to 3) and VIP (Day 1 only): genotype * condition interaction: Day 1, F(1,1447)=2.240, p=0.1347 ; Day 2, F(1,1293)=49.99, p<0.0001 ; Day 3, F(1,1227)=117.6, p<0.0001; α=0.05. **(b)** The activity levels across 10 minutes in the curved track. O-LM were more active than VIP: cell type effect: Day 1, F(1,1460)=55.37, p<0.0001 ; Day 2, F(1,1400)=362.2, p<0.0001 ; Day 3, F(1,817)=1=268.6, p<0.0001; α=0.05. **Statistics: (a-b)** 3-way ANOVA with LSD multiple comparisons post hoc test. Differences between days, for the same genotype, same measure &&&& p<0.0001. Differences between groups on the same day * p<0.05, ** p<0.01, *** p<0.001 and, **** p<0.0001. Data are shown as Mean ± SEM.

**Extended Data Fig. 5.**
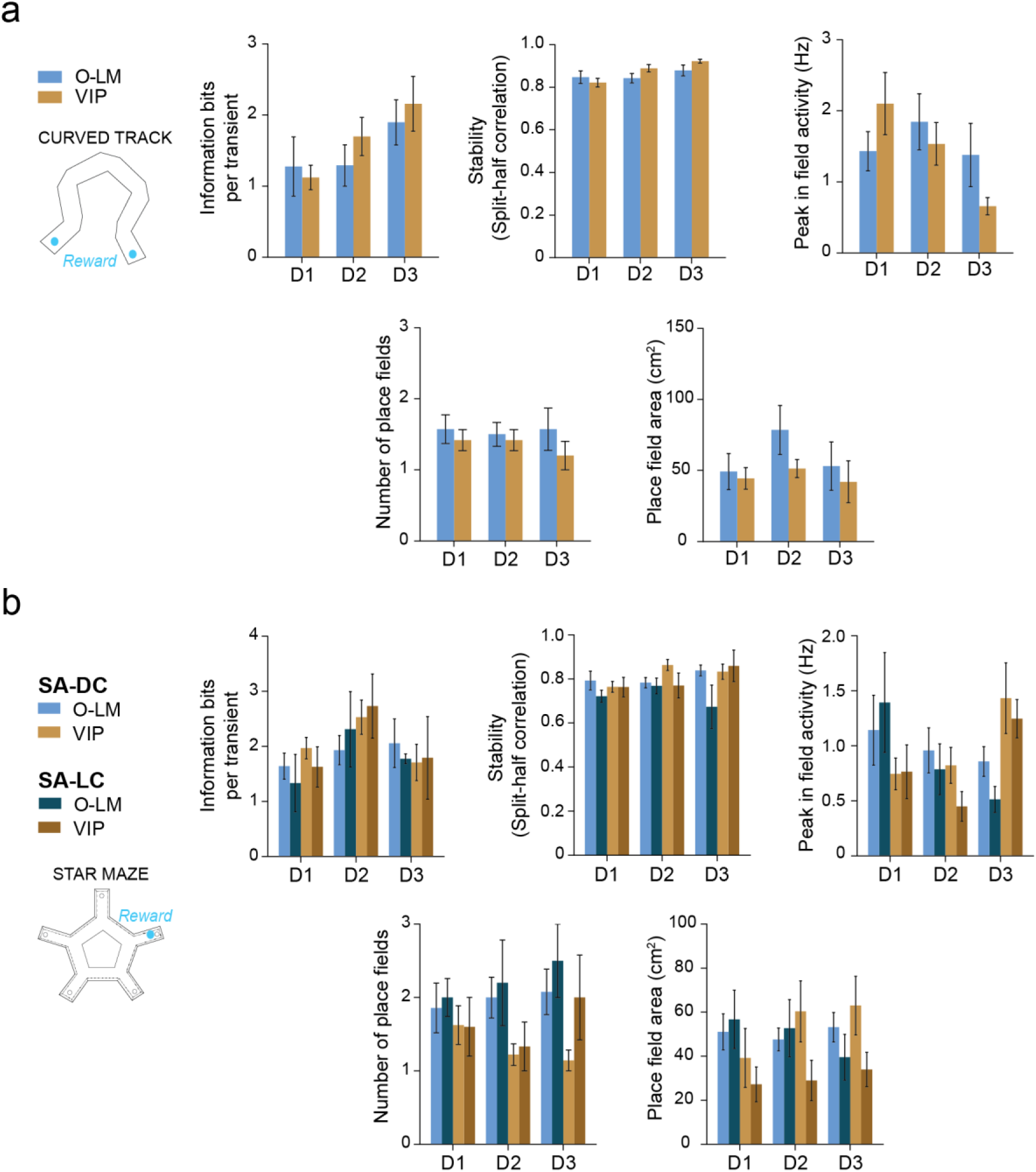
Interneurons’ place field metrics were not affected by the cell type nor the behavioral procedure. **(a)** Place cell metrics in the curved track. Spatial information bits per calcium transient did not differ between O-LM and VIP place cells. Place cell stability did not differ between O-LM and VIP, but increased with familiarization (session effect: F(2,52)=3.396, p=0.0411). Place cells’ peak firing rate in the field did not change significantly. Neither the number of place fields per cell nor the average field size differ between O-LM and VIP. **(b)** Place cell metrics in the Star Maze. For both SA-DC and SA-LC conditions, spatial information bits per calcium transient did not differ between O-LM and VIP place cells. Place cell stability was slightly reduced in the SA-LC condition (condition effect: F(1,73)=5.187, p=0.0257). Place cells’ peak firing rate within place field decreased across session in O-LM but increased in VIP, independently of the condition (cell type * session interaction: F(2,73)=3.738, p=0.0285). The number of place fields per cell was higher in O-LM compared with VIP independently of the condition or the session (cell type effect: F(1,69)=7.408, p=0.0082). Average place field size did not differ significantly between O-LM and VIP and across conditions.

**Extended Data Fig. 6.**
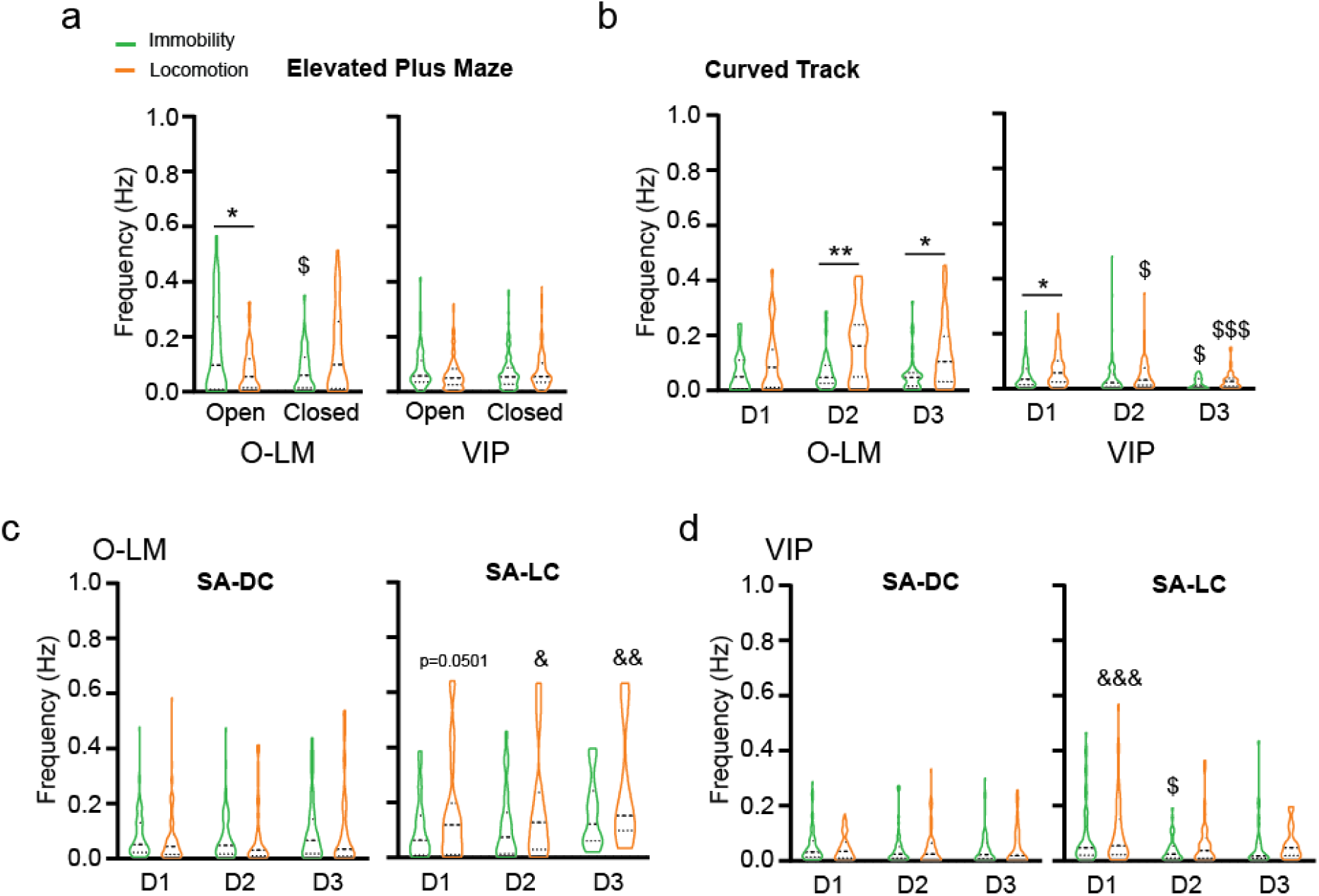
Change in O-LM and VIP firing rate during locomotion and immobility. Mean cell frequencies recorded for both O-LM and VIP interneurons, during locomotion (orange) and immobility (green) in (**a**) the EPM (**b**) the curved track and (**c-d**) the two Star Maze conditions. (**a**) in the EPM, O-LM were more active during immobility in open arms and more active during locomotion in closed arms in contrast to VIP (Locomotion state * cell type * arm type interaction: F(1,412)=16.04; p<0.0001). (**b**) In the curved track, on Day 1 interneurons activity was independently influenced by the cell type and the locomotion state (cell type effect: F(1,146)=4.553, p=0.0345; locomotion state effect: F(1,146)=24.90, p<0.0001; cell type * locomotion state interaction: F(1,146)=2.241, p=0.1365). On Day 2 and 3, only O-LM increased their firing rate during locomotion (Day 2: cell type * locomotion state interaction: F(1,140)=35.51, p<0.0001 and Day 3 : cell type * locomotion state interaction: F(1,83)=20.67, p<0.0001). (**c-d**) In the Star maze, O-LM, but not VIP-pyr, firing rate was consistently higher during locomotion in the SA-LC condition (cell type * locomotion state * condition interaction: Day 1: F(1,230)=6.405, p=0.0120; Day 2: F(1,199)=11.88, p=0.0007; Day 3: F(1,195)=8.181, p=0.0047). **Statistics:** (**a**) differ between arms for the same behavioral state $ p<0.05. (**b,d**) differ from the same behavioral state recorded on Day 1 $ p<0.05, $$ p<0.01, $$$ p<0.001 and $$$$ p<0.0001. (**a-d**) Differ between behavioral states on the training session * p<0.05, ** p<0.01. (**c-d**) Differ from the same behavioral state, same training session recorded in the SA-DC & p<0.05, && p<0.01 and &&& p<0.001. Data are shown as Mean ± SEM.

**Extended Data Fig. 7.**
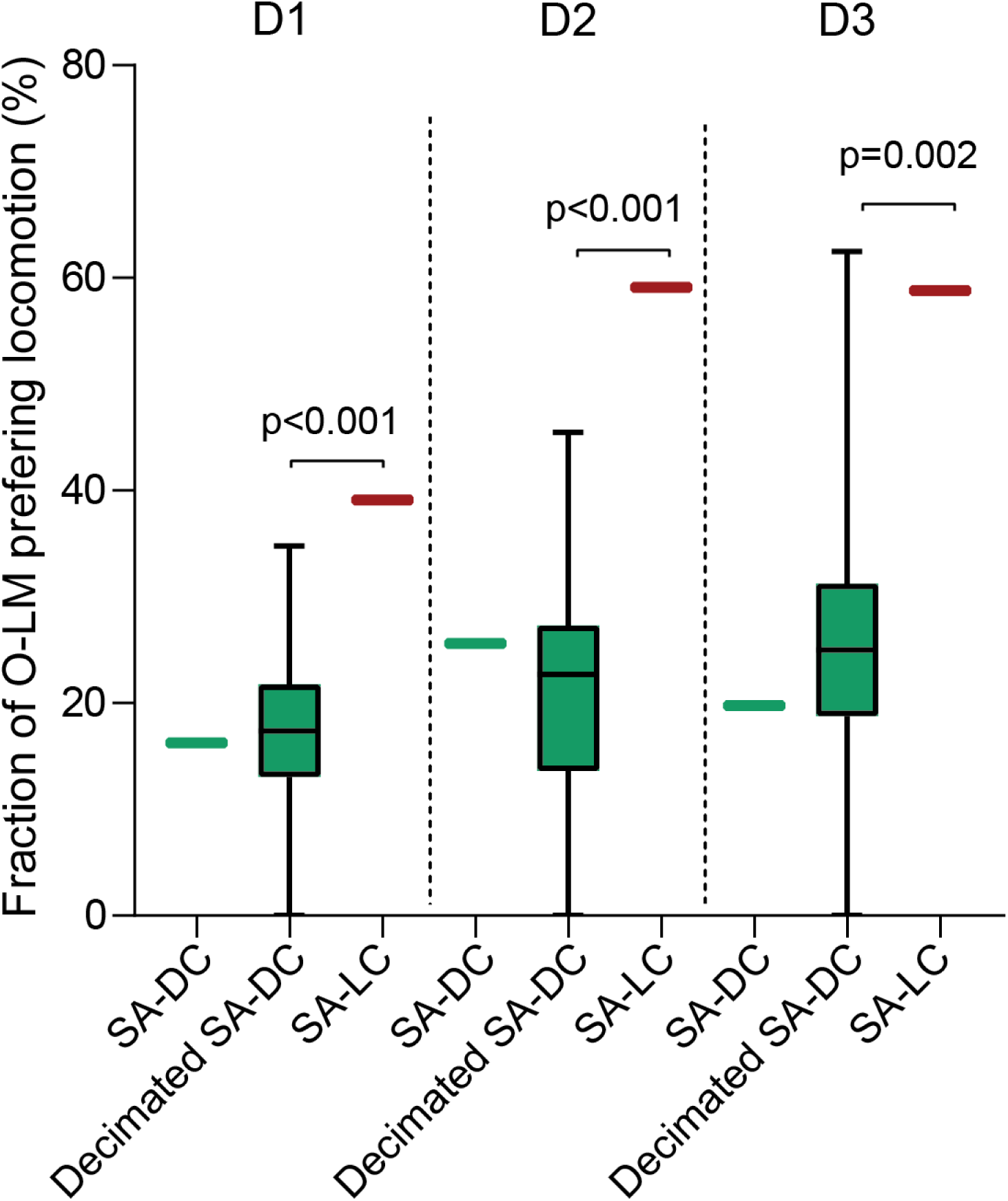
Observed probability of replicating the O-LM preferential activation during locomotion displayed in the SA-LC condition by decimating the SA-DC condition dataset. Decimation of SA-DC O-LM failed to recapitulate SA-LC O-LM locomotion preference, but rather replicate fraction of O-LM preferring locomotion observed in the original SA-DC dataset. Briefly, for each of the 1000 iterations of decimation on the SA-DC dataset, the fraction of O-LM preferring locomotion was computed using time-shuffling analysis (whisker box), the original fraction of locomotion preferring O-LM from SA-DC condition is denoted by the green bar on the left of the whisker box, and the fraction of locomotion preferring O-LM from the SA-LC condition is demoted by the red bar on the right of each whisker box. Probabilities on the graph are the observed number of decimation iterations that generated an equal or superior fraction of locomotion-preferring O-LM than the one observed in the SA-LC dataset. One-sample t-test vs. SA-LC fraction : D1 t(999)=112.7, p<0.0001; D2 t(999)=162.6, p<0.0001; D3 t(999)=78.54, p<0.0001.

**Extended Data Fig. 8.**
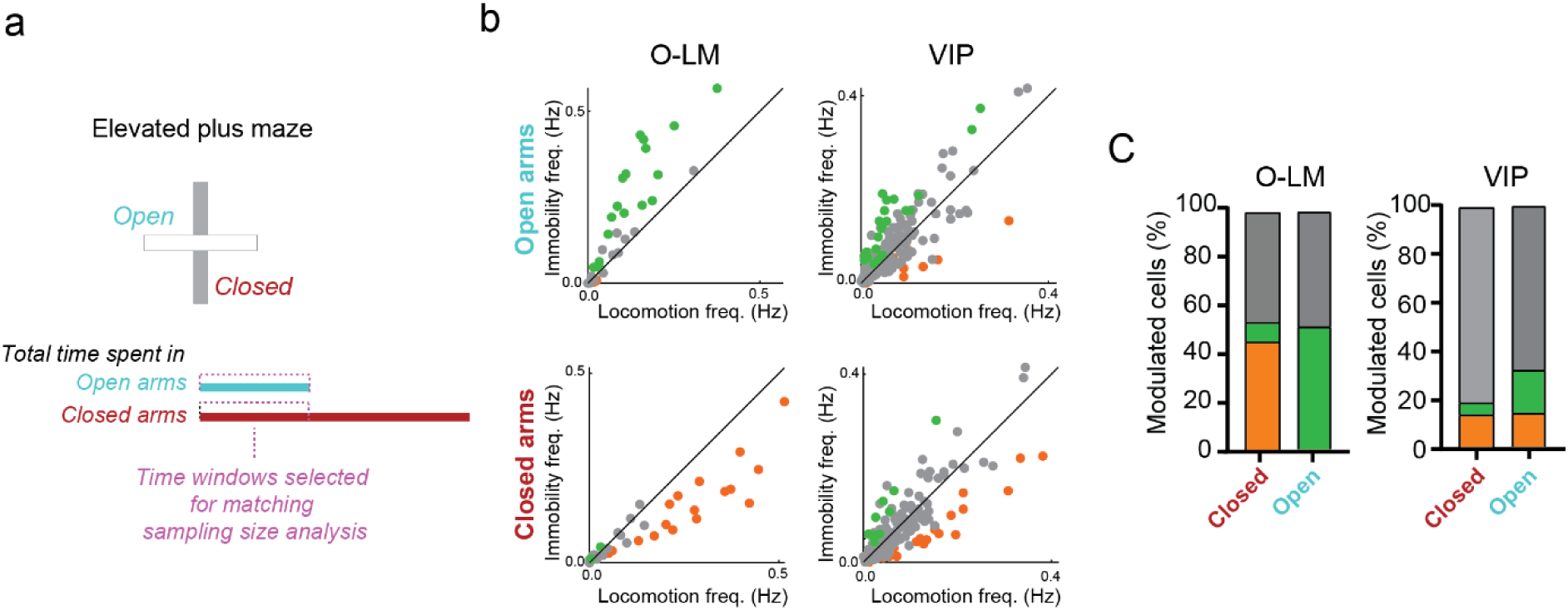
Time-matching analysis of EPM calcium activity confirmed that O-LM interneurons are preferentially tuned to locomotion in closed arms and immobility in open arms. (**a**) As mice spent more time in the closed arms than in open arms, we checked that the differences reported on fig. 5 were not due to this difference in sampling size. Briefly, for each mouse we re-run the analysis restricting the closed arm dataset to match the duration of open arms exploration. (**b**) Scatter-plot for EPM open arms (Top) and closed arms (Bottom) locomotion vs. immobility activity. Each dot represents an individual interneuron. O-LM preference shifted from a preferential activation during locomotion in closed arms to immobility in open arms whereas VIP were split between locomotion-preferring and immobility-preferring cells in both arms. (**c**) Corresponding fractions of O-LM and VIP cells significantly modulated by locomotion (orange), immobility (green) or with no significant preference (gray) as determined by time-shuffling analysis. Left: the proportion of O-LM interneurons that prefer locomotion was higher in closed arms (45% vs. 0%, x2 = 55.5125, p<0.0001) whereas the proportion of O-LM preferring immobility was higher in open arms (48% vs. 10%, x2 = 33.2443, p<0.0001). Right: the proportion of VIP interneurons that prefer locomotion was similar in closed and open arms (15% vs. 16%, x2 = 0, p=1) whereas the proportion of VIP preferring immobility was higher in open arms (6% vs. 16%, x2 = 4.1369, p=0.0420).

**Extended Data Fig. 9.**
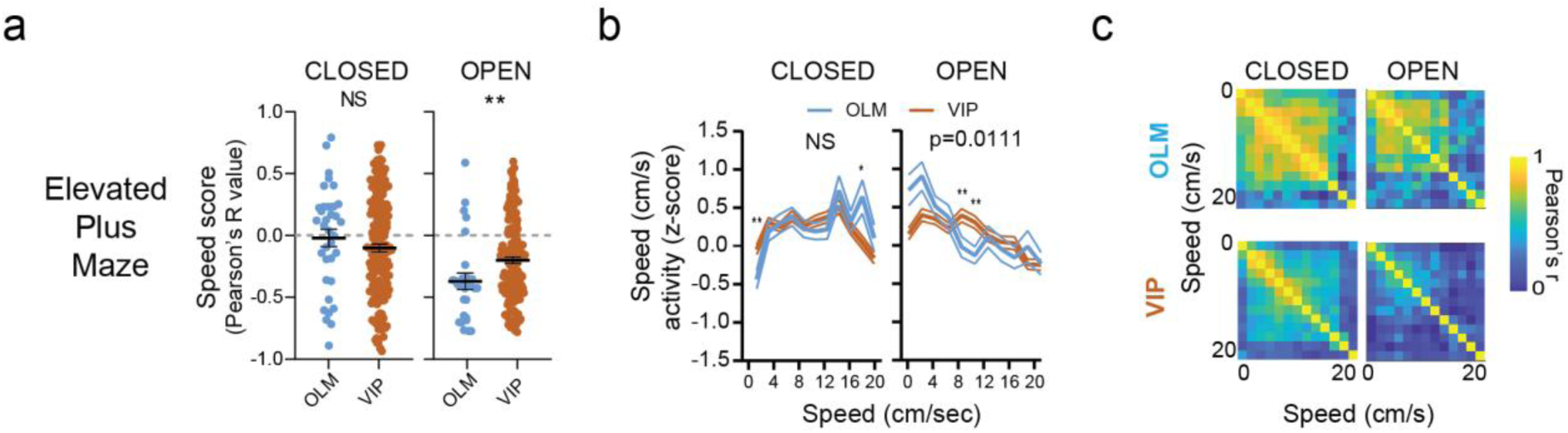
Time-matching analysis of EPM calcium activity confirmed the O-LM bi-modal speed coding properties in closed vs. open arms. As mice spent more time in the closed arms than in open arms, we checked that the differences reported on fig. 6 were not due to this difference in sampling size. Briefly, for each mouse we re-run the speed analysis while downsampling the closed arm dataset to match the duration of open arms exploration. (**a**) re-analysis with time-matched closed arm data confirms that O-LM cells speed scores were lower than VIP cells in open arms but similar in closed arms (open: t(188)=2.411, p=0.0169 ; closed: t(200)=1.052, p=0.2942 ; α=0.05). (**b**) re-analysis with time-matched closed arm data confirms that in contrast to VIP, O-LM interneurons respond differentially to increasing speed in each arm type, increasing their activity with speed in closed arms whereas decreasing their activity with increasing speed in open arms (speed * arm * cell type interaction: F(10,1862) = 3.103, p=0.0006; α=0.05). (**c**) PV analysis on time-matched dataset confirms that VIP cells maintained a population coding of the speed in both arms whereas O-LM cells may display a rate coding of the speed, particularly in the closed arm. **Statistics:** (**a**) two-tailed unpaired t test: ** p<0.01, *** p<0.001. (**b**) On graph: 2-way ANOVA with Bonferroni multiple comparisons post hoc test; difference between SM cell type: * p<0.05, ** p<0.01, *** p<0.001. In the legend: 3-way ANOVA. Data are shown as Mean ± SEM.

**Extended Data Fig. 10.**
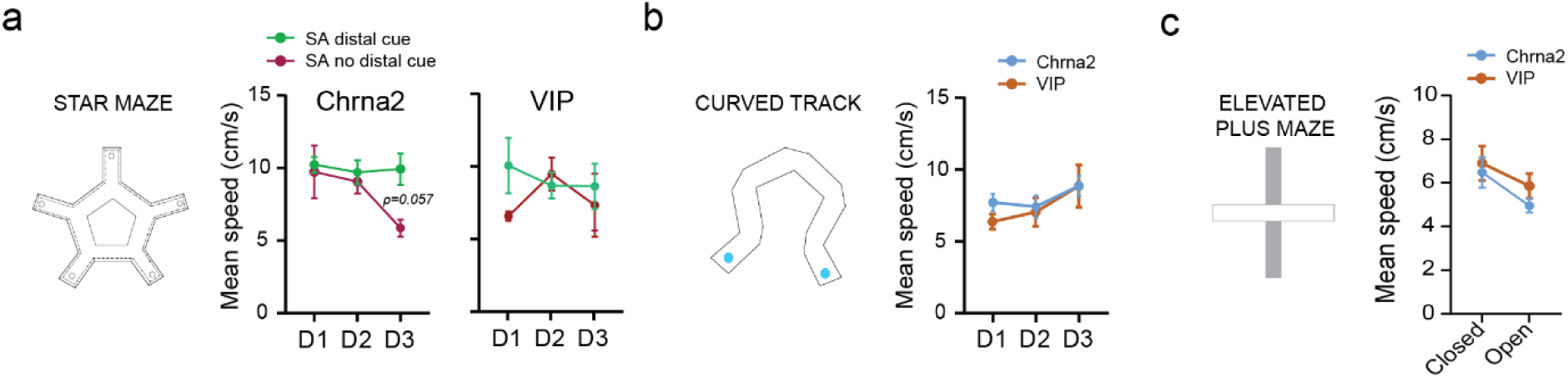
Running speed during navigation. Chrna2-cre and VIP-cre mice’s speed performances for **(a)** Star maze conditions SA-DC in green and SA-LC in magenta, **(b)** Curved track and **(c)** Elevated plus maze tasks. **(a-c)** No significant differences were found between genotypes for all behavioral paradigms employed. **(a-b)** No significant influence of training was recorded on animals’ speed performances. **(b-c)** speed performances from Chrna2-cre mice (light blue) and VIP animals (brown).

**Extended Data Fig. 11.**
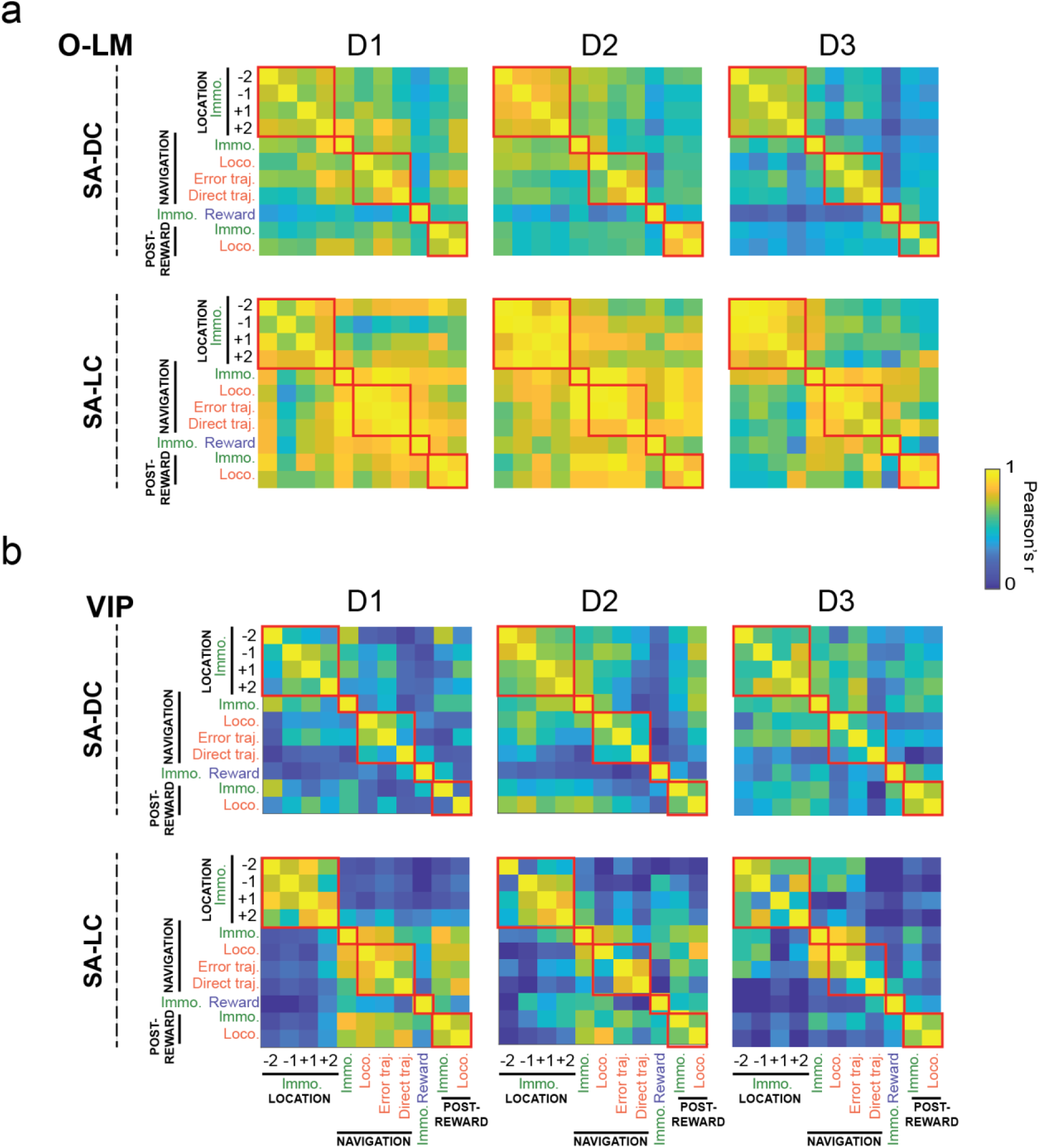
Orthogonality of O-LM and VIP population activity using pseudo-population vectors of the different behavioral phases. **(a-b)** Pseudo-population vector correlation matrices for behavioral events in the SA-DC (top) and SA-LC (bottom) procedures across learning days. The firing rates for all recorded interneurons in a given state were correlated across each pairwise behavioral state combination. High correlation values (yellow) indicate similar population activity for the two states. **(a)** In the SA-DC condition, O-LM PV for behavioral events became more orthogonal with time in comparison to the SA-LC condition, suggesting that O-LM interneuron sub-networks specialize for different behavioral events with time only when mice relied on a distal frame of reference. **(b)** VIP interneuron behavioral event PV maintains a degree of orthogonality in their activation in both Star maze conditions, suggesting a lack of sensitivity to the frame of reference used.

**Extended Data Fig. 12.**
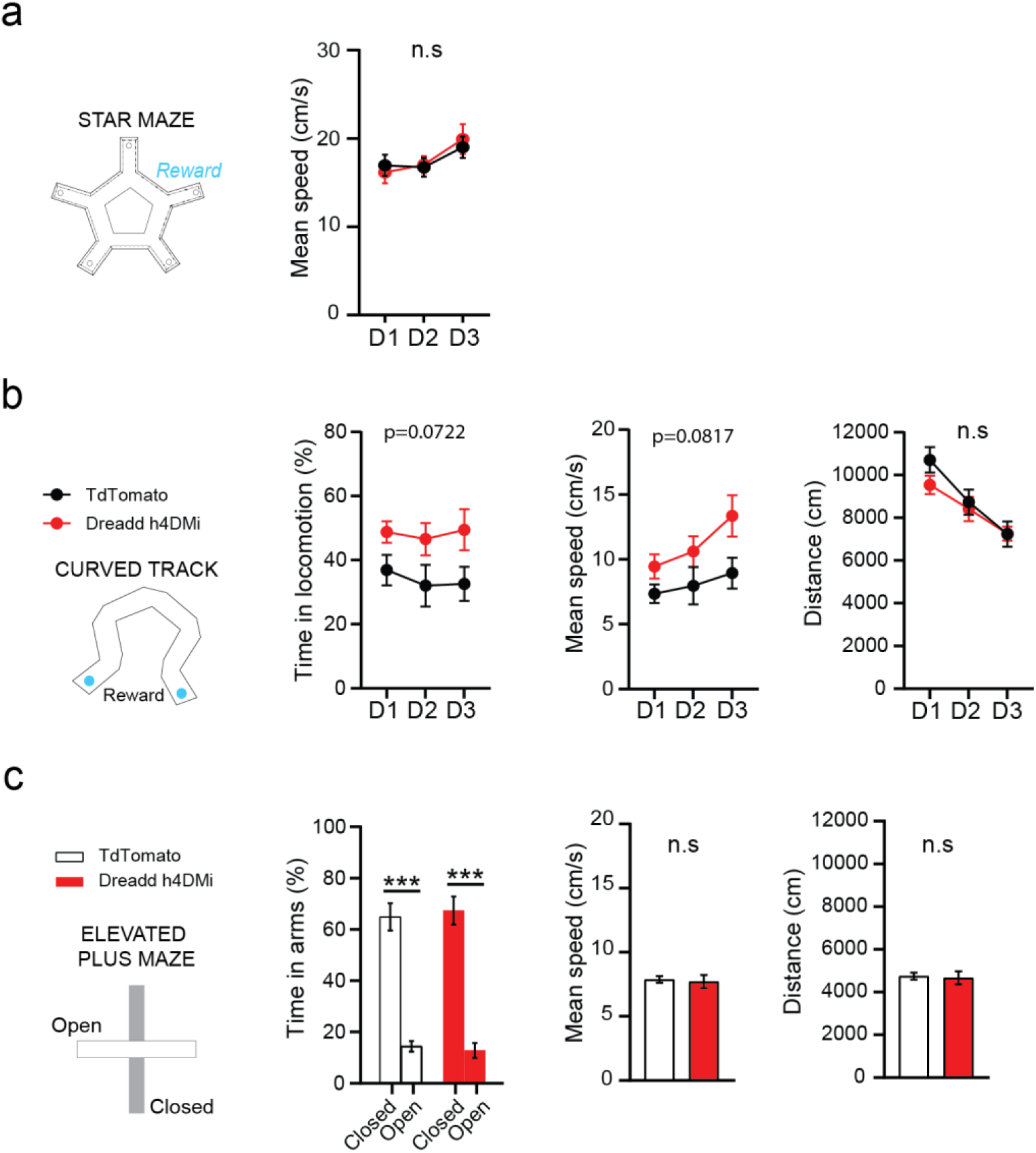
Chemogenetic inhibition of dCA1 O-LM interneurons does not alter anxiety nor locomotion-associated behaviors. (**a**) Speed performances were unchanged by Dreadd h4DMi inhibition of dCA1 OLM interneurons across the training session (D1-D3) in the SA-DC condition of the star maze. (**b**) No significant changes were observed in the percentage of time spent in locomotion (left), the averaged speed of displacements (middle) nor in the total distance traveled (right) in the curved track paradigm. (**c**) No significant changes were detected in the percentage of time spent into the closed *vs*. open arms (left). Both control and experimental animals showed a preference for closed areas of the maze. Mean speed (middle) and total distance (right) remain unchanged between experimental conditions (right). Two-way Repeated ANOVA for the effect of C21, arms and their interaction: C21: F(1,10) = 0.03621, p = 0.8529; arms: F(1,10) = 94.79, p < 0.0001; interaction: F(1,10) = 0.1408, p = 0.7154. **Statistics:** Panels **(c)** were analyzed via two-way repeated ANOVA followed by Bonferroni’s multiple comparisons *** p < 0.001). Data are presented as Mean ± SEM.

